# Stomatal closure in maize is mediated by subsidiary cells and the PAN2 receptor

**DOI:** 10.1101/2023.04.29.538816

**Authors:** Le Liu, M. Arif Ashraf, Taylor Morrow, Michelle Facette

**Affiliations:** Department of Biology, University of Massachusetts Amherst, Amherst, MA, USA 01003; Department of Biology, Howard University, Washington, DC, USA 20059

**Keywords:** confocal microscopy, corn, gas exchange, guard cell, maize, stomata, subsidiary cells

## Abstract

Stomata are epidermal pores that facilitate plant gas exchange. Grasses have fast stomatal movements, likely due to their dumbbell-shaped guard cells and lateral subsidiary cells. Subsidiary cells reciprocally exchange water and ions with guard cells. However, the relative contribution of subsidiary cells during stomatal closure is unresolved. We compared stomatal gas exchange and stomatal aperture dynamics in wild type and *pan1*, *pan2*, and *pan1;pan2 Zea mays (L.)* (maize) mutants, which have varying percentages of aberrantly formed subsidiary cells. Stomata with 1 or 2 defective subsidiary cells cannot close properly, indicating that subsidiary cells are essential for stomatal function. Even though the percentage of aberrant stomata is similar in *pan1* and *pan2, pan2* showed a more severe defect in stomatal closure. In *pan1*, only stomata with abnormal subsidiary cells fail to close normally. In *pan2*, all stomata have stomatal closure defects, indicating that PAN2 has an additional role in stomatal closure. Maize *Pan2* is orthologous to Arabidopsis *GHR1,* which is also required for stomatal closure. PAN2 acts downstream of Ca^2+^ in maize to promote stomatal closure. This is in contrast to GHR1, which acts upstream of Ca^2+^, and suggests the pathways could be differently wired.

## Introduction

Stomata are small pores on the leaf surface of land plants that facilitate gas exchange between the plant and the atmosphere. Acquisition of stomata and leaf cuticles was critical for plants to colonize the fluctuating terrestrial environment (Hetherington & Woodward, 2003). Open stomatal pores allow the absorption of carbon dioxide into the plant for photosynthesis, while simultaneously allowing oxygen and water vapor to diffuse out. Stomata are responsible for ∼80-95% of plant transpiration, depending on species and leaf water potential (Hanson *et al*., 2016). Plants regulate stomatal opening and closing (i.e., regulate stomatal aperture) in response to exogenous and endogenous signals to balance water loss and carbon gain. Exogenous signals such as light, vapor pressure deficit (VPD), and CO_2_ concentration regulate stomatal aperture (Eyland *et al*., 2021). Endogenous signals that regulate stomatal aperture include phytohormones like abscisic acid (ABA), hydrogen peroxide (H_2_O_2_), nitric oxide (NO), and other mesophyll-derived metabolites (Suhita *et al*., 2004; Munemasa *et al*., 2015; Lawson & Matthews, 2020). The time it takes for stomata to open or close in response to light cues is much greater than how long it takes for photosynthetic light reactions to be activated or stopped (Pearcy, 1990; Lawson & Blatt, 2014). Therefore, photosynthetic capacity is often dictated by the rate of stomatal movement when light intensity is not limiting. The kinetics of stomatal conductance (*g_s_)* influences plant growth and water use efficiency in the field (Lawson and Blatt, 2014). It has been hypothesized that plants with faster stomatal kinetics have advantages in water use and photosynthesis efficiency in variable environments since steady-state values of *g_s_* and assimilation under fluctuating conditions can be achieved rapidly (Lawson & Blatt, 2014; Vialet-Chabrand *et al*., 2017; Eyland *et al*., 2021; Long *et al*., 2022).

The relationship between guard cells and their neighbors has been documented, including the “mechanical advantage” that adjacent cells have over guard cells (DeMichele & Sharpe, 1973). Laser ablation of cells adjacent to guard cells leads to an increase in stomatal aperture in *Arabidopsis thaliana (L.)* (Davaasuren *et al*., 2022). Graminoid (grass) stomata have rapid stomatal responses relative to other plants (Raschke & Fellows, 1971; Grantz & Assmann, 1991; Franks & Farquhar, 2007; Vico *et al*., 2011; McAusland *et al*., 2016; Lawson & Vialet-Chabrand, 2019). The unique dumbbell-shaped guard cells and laterally associated subsidiary cells likely contribute to the fast graminoid stomatal movements (Hetherington & Woodward, 2003; Franks & Farquhar, 2007; Gray *et al*., 2020; Nunes *et al*., 2020). The reciprocal exchange of potassium and chloride between subsidiary cells and guard cells was first shown in *Zea mays (L.)* (maize) by Raschke and Fellows. This “see-saw” model of a reciprocal flux of water and ions was further corroborated (Franks & Farquhar, 2007) and has been shown in barley (Schäfer *et al*., 2018). Finite element modeling of guard cell and subsidiary cell pressures coupled with morphometric analyses suggest that subsidiary cells are important to enhance pore closure in grasses (Durney *et al*., 2023). Genetic evidence supporting the importance of subsidiary cells was first demonstrated via the *Brachypodium distachyon (L.)* mutant *sid*, and shortly thereafter by the maize mutant *bzu2* (Raissig *et al*., 2017; Wang *et al*., 2019a). The mutated genes that cause the *sid* and *bzu2* phenotypes in *B. distachyon* and maize are both orthologous to the *A. thaliana* bHLH transcription factor *AtMUTE,* therefore the genes are referred to as *BdMUTE* and *ZmMute.* In *A. thaliana*, MUTE is important for the specification of guard cell fate (Pillitteri & Torii, 2007). In *B. distachyon* and *Z. mays*, loss of MUTE results in a complete loss of subsidiary cells. Both grass mutants have defects in stomatal closure. However, *sid* and *bzu2* also have defective guard cell divisions to various extents, which result from aberrant or failed division of the guard cell progenitor. *B. distachyon sid* has a mild guard cell phenotype, while maize *bzu2* has an especially severe guard cell phenotype that leads to seedling lethality (Raissig *et al*., 2017; Wang *et al*., 2019a). Therefore, to assess the role of subsidiary cells, analysis of additional mutants is warranted.

The mechanism of turgor-driven movement of guard cells has been well-researched using *A. thaliana* as a model (Jezek & Blatt, 2017; Lawson & Matthews, 2020; Hsu *et al*., 2021). In *A. thaliana*, signals that trigger stomatal closure are typically mediated by ABA and core guard cell-specific kinases (Nunes *et al*., 2020; Hsu *et al*., 2021). Stomatal closure results from the activation of ion channels and transporters, including the slow-type anion channel (SLAC1), which allows the efflux of K+ and then leads to water efflux and causes turgor decrease in the guard cells, which in turn leads to closure of the stomatal pore (Negi *et al*., 2008; Vahisalu *et al*., 2008; Geiger *et al*., 2009; Munemasa *et al*., 2015). In Arabidopsis, SLAC1 is potentially activated by the leucine-rich repeat receptor-like pseudokinase (LRR-RLK) GUARD CELL HYDROGEN PEROXIDE-RESISANT1 (GHR1), in addition to other signaling components such as the OST1 kinase (Hua *et al*., 2012). Despite being a pseudokinase, several independent lines of evidence suggest that GHR1 physically interacts with SLAC1, and that GHR1 might activate SLAC1 (Hua et al., 2012; Sierla et al., 2018). Physical interactions between GHR1 and SLAC1 have been demonstrated by bimolecular fluorescence complementation (BiFC), firefly luciferase complementation, and coimmunoprecipitation (Hua *et al*., 2012; Sierla *et al*., 2018). Using split-GFP constructs, GHR1 can activate SLAC1 in *Xenopus oocytes* (Sierla *et al*., 2018). Both *ghr1* and *slac1* have stomatal closure defects in response to various stimuli, and *ghr1slac1* shows the same phenotype as *slac1* (suggesting the same pathway).

The increase of cytosolic H_2_O_2_ and Ca^2+^ of guard cells acts as a molecular switch that modulates the activity of various ion channels, transporters, and enzymes involved in stomatal movement, leading to the closure of stomata and reducing water loss in *A. thaliana* plants (Hua *et al*., 2012; Sierla *et al*., 2018; Hsu *et al*., 2021). Stomatal closure is stimulated by ABA, H_2_O_2_ and Ca^2+^ in maize just as it is in *A. thaliana*. Notably, more cytosolic H_2_O_2_ and Ca^2+^ accumulates in subsidiary cells than in guard cells during maize stomatal closure (Yao *et al*., 2013; Zhang *et al*., 2020).This raises the question of how subsidiary cell turgor is regulated. Subsidiary cells have opposite ion fluxes and volume changes (and by inference, opposite turgor) to guard cells (Nunes *et al*., 2020), but how does this occur? Are there different channels and/or regulatory proteins in subsidiary cells, or are the same mechanisms in guard cells employed in subsidiary cells? Are guard cells primarily regulated, and subsidiary cells merely responsive to the changes in the adjacent guard cells, perhaps accumulating potassium and calcium the guard cells export into the apoplast? Previous analyses have identified proteins shared between grass subsidiary and guard cells, as well as those that are unique to each cell type (Majore *et al*., 2002; Mumm *et al*., 2011; Nguyen *et al*., 2017; Wang *et al*., 2019b; Sun *et al*., 2022). Nonetheless, whether the two cell types have unique signaling mechanisms, and what if any coordination exists between subsidiary cells and guard cells is still unclear.

To understand the relative roles of subsidiary cells to the stomatal closure, we assayed stomatal function using previously characterized maize mutants which have defects in subsidiary cell formation. The *pan1* and *pan2* mutants have subsidiary cell-specific defects in division plane orientation, leading to abnormally shaped subsidiary cells that are often not correctly specified (Gallagher & Smith, 1999; Zhang *et al*., 2012). *Pan1* and *Pan2* both encode catalytically inactive leucine-rich repeat receptor-like pseudokinases (LRR-RLKs). Despite their similar names, and the fact they encode the same type of protein, *pan1* and *pan2* are not paralogs. They share a gene name because they were discovered in the same screen. Maize *Pan2* is orthologous to Arabidopsis *GHR1* (Man et al., 2020). We found that stomatal complexes with abnormal subsidiary cells impaired stomatal closure, indicating an essential role for subsidiary cells. Moreover, *Pan2*, just as its one-to-one *A. thaliana* orthologue *GHR1* does, plays an essential role in the grass stomatal closure. Overall, our study provides insights into the relative roles of subsidiary cells and guard cells in stomatal closure and highlights the importance of PAN2 in the grass stomatal closure.

## Materials and Methods

### Plant materials and Growth Conditions

*Zea mays (L.)* (maize) mutants *pan1-ems* (Cartwright *et al*., 2009), *pan2-O* and *pan2-2* and *pan1-Mu;pan2-O* (Zhang *et al*., 2012) have been previously described. All mutants were backcrossed at least five times into the B73 wild-type background. B73 inbred lines were used as controls in all experiments. PAN1-YFP and PAN2-YFP transgenic plants were described previously (Humphries *et al*., 2011; Zhang *et al*., 2012). These are genomic constructs with the native promoter and terminator. All the plants used in this study were grown in a greenhouse between 20°C and 29°C with supplemental LED light (Fluence VYPR series) (16-hour day length). Plants were grown in Pro-mix Professional soil supplemented with Peters Excel 15-5-15 CalMag fertilizer and chelated liquid iron (Southern Ag) as needed.

### Gas exchange measurements

Stomatal conductance measurements were performed using a CIRAS3 gas analyzer equipped with a PCL3 leaf cuvette (PP Systems). For all measurements, healthy, fully expanded leaves were used. Briefly, the leaf cuvette was clamped onto the leaves and equilibrated for at least 30 mins to stabilize the initial stomatal conductance at 1000 PPFD. The CO_2_ concentration was held constant at 410 parts per million (ppm); H_2_0 reference was set to 80%; the cuvette flow at 300 cc min^-1^. Stomatal conductance (*g_s_*) and assimilation (A) rates were measured every 5 seconds during the entire treatment period. To measure stomatal closure rates, lights were turned off (PPFD=0) after the initial conductance became stable. For CO_2_ responses, after the initial stomatal conductance was steady, and CO_2_ concentration was shifted from 410 ppm to 2,000 ppm. Relative *g_s_* was computed for each individual measured plant by normalizing *g_s_* to the initial steady state *g_s_* value observed. Data analysis was performed in R and JMP® 16.1. The data presented are the mean ± standard error of at least five leaves per treatment.

### Stomatal phenotyping

Epidermal glue impressions were performed on fully expanded leaf 3, 5 or 8 according to (Allsman *et al*., 2019). The percentage of stomata with one or two abnormal subsidiary cells, and stomatal density quantification is based on glue impressions, which were imaged using a Nikon SMC800N dissecting microscope.

Morphology of guard and subsidiary cells and guard cell length was examined via confocal microscopy (described below). Plants were fixed in formaldehyde-alcohol-acetic acid (FAA) and then stained with 0.1% Direct Red and 0.1% Calcofluor.

### Stomatal aperture assays

All the leaves we included in the stomatal aperture experiment were healthy and fully expanded. Maize leaf blade segments from leaf 5 or leaf 8 were floated in opening buffer (10 mM KCl, 100μM CaCl_2,_ and 10 mM MES/Tris, pH 5.6) (Zhang *et al*., 2020) in the absence or presence of 1 mM ABA (Rodriguez & Davies, 1982) under light (500-700 PPFD). For Ca^2+^ treatment experiments, the epidermis was peeled from fully expanded L4 and immediately floated in opening buffer supplemented (or not) with an additional 10 mM CaCl_2_ under light. Leaves were treated for 2 hours; then, the leaves were stained using 1mM Propidium iodide (PI) for 15 mins (whole leaf piece) or 1 min for peels. The stomatal aperture from live (unfixed) leaves were immediately observed under a confocal microscope (described below) and measured using FIJI software (Schindelin *et al*., 2012).

### Gene Expression Analysis by qRT-PCR

RNA was extracted from the division zone and different leaf samples from greenhouse-grown maize plants using RNeasy® Plant Mini Kit (Qiagen). Extracted RNA concentrations were quantified using a NanoDrop (Thermo Scientific). Each RNA concentration was normalized with RNase-free water, and 500 ng RNA was used to synthesize cDNA using ProtoScript® II Reverse Transcriptase (NEB). Quantitative PCR reactions were performed using StepOnePlus™ Real-Time PCR System (Applied Biosystems) and PowerUpTM SYBR^TM^ Green Master Mix (Applied Biosystems). The reaction was performed as per the manufacturer’s instructions. For quantification of gene expression, we used the 2^−ΔΔCT^ (cycle threshold) method with normalization to the UBCP expression. Data were obtained from three biological replicates (independent plants) and two technical replicates. Primers used in this study are listed in Supplemental Table S1.

### Plasmolysis

Leaf blade segments from fully expanded maize leaf 4 expressing PAN2-YFP were submerged into 500 mM mannitol for 10 mins. Segments were then mounted in 500 mM mannitol and immediately observed using confocal microscopy.

### Confocal Microscopy

Plant tissues were observed using a custom-built spinning disc confocal unit (3i) equipped with an inverted fluorescence microscope (IX83-ZDC, Olympus) CSU-W1 spinning disc with 50-micron pinholes (Yokogawa), a Mesa Illumination Enhancement 7 Field Flattening unit (3i), an Andor iXon Life 888 EMCCD camera, a UPLANSAPO60XS2 (NA = 1.20, Olympus) or UPLANSAPO40X (NA = 1.25, Olympus) Silicone Oil Immersion objective and 4 laser stacks with TTL controller (3i). For YFP imaging of transgenic maize plants, the 514 nm laser and 542/27 emission filter (YFP) were used. The 568 laser with 618/50 emission filters was used for propidium iodide and Direct Red staining experiments. Image processing was performed using Image FIJI and Adobe Photoshop version 8.0. The YFP fluorescence intensity was qualitatively measured using FIJI. Max projections of six slices of 16-bit images taken on the same day with the same acquisition settings were used. Sibling plants without YFP fluorescence were used as controls.

### Accession Numbers

Gene and protein sequences for maize and Arabidopsis can be found at MaizeGDB (www.maizegdb.org) and TAIR (https://www.arabidopsis.org/index.jsp), respectively. Accession numbers for maize Version 5.0 of the V73 genome are: PAN1 (Zm00001eb034900), and PAN2 (Zm00001eb117610).

### Statistical Analyses

Non-parametric comparisons for each pair using Wilcoxon Methods were performed to compare stomatal aperture, and a Bonferroni multiple testing correction was applied to generate p-adjusted (p-adj). Summary statistics for stomatal aperture are listed in Supplemental Table S2 and results of statistical tests are in Supplemental Table S3. Tukey Kramer t-test was used to compare the percentage of abnormal stomata and Student’s t-test was used for qPCR value. Raw CT value used for statistical test is in Supplemental Table 4, and the results of the

T-tests are listed in Supplemental Table 5. All statistical tests were performed in JMP® 16.1.

## Results

### The percentage of abnormal stomatal complexes varies across different genotypes and leaves

What is the relative contribution of subsidiary cells in grass stomatal function? Are subsidiary cells a passive reservoir that accumulate ions and water because of their proximal location to guard cells, or do they actively promote stomatal function? PANGLOSS1 (PAN1) and PAN2 were previously identified as catalytically inactive LRR-RLKs in maize that are polarized prior to the formative divisions that generate subsidiary cells during stomatal development (Cartwright *et al*., 2009; Zhang *et al*., 2012). Both *pan1-ems* and *pan2-2 are* null mutants that have incomplete penetrance, where only a portion of subsidiary cells are abnormal in the single mutants (Cartwright *et al*., 2009; Zhang *et al*., 2012; Figure 1). The percentage of abnormal subsidiary cells varies by leaf number in *pan1* and *pan2* (Figure 1). Leaves that emerge first have a higher frequency of subsidiary cell defects, while later-emerging leaves have fewer defective cells. Previous analyses suggest that morphologically correct subsidiary cells are likely correctly specified (Gallagher & Smith, 1999). We are unable to propagate the more severe *pan1-ems;pan2-2* double mutants, so the weaker alleles of *pan1-mu* and *pan2-O* were used to generate *pan1-mu;pan2-O.* The *pan1;pan2* double mutant has a synergistic phenotype, with more than double the frequency of abnormal subsidiary cells than either of the single mutants (Zhang *et al*., 2012; Figure 1).

**Figure 1:**
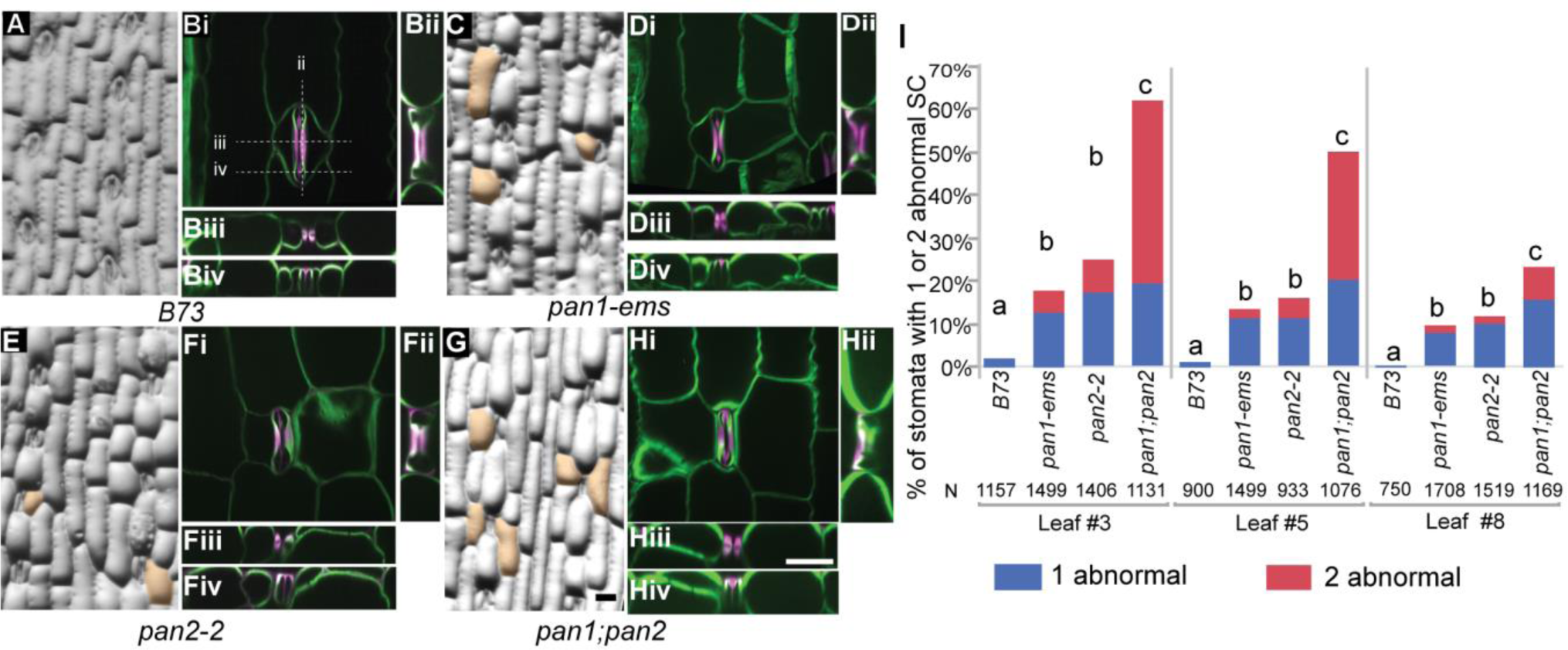
Mature subsidiary cells are abnormal in *pan1*, *pan2,* and *pan1;pan2* mutants. Epidermal glue impressions (A, C, E, G) and confocal images (B, D, F, H) of wild type (A, B), *pan1* (C,D), *pan2* (E,F), and *pan1;pan2* (G,H). Abnormal subsidiaries are false-colored orange in the glue impressions. The confocal images show individual stomates stained with direct red (magenta) and calcofluor (green) and include reconstructed orthogonal views in the x (ii, right of image) and y (iii & iv, below image) planes. (E) Percentage of stomatal complexes with abnormal subsidiary cells in leaf 3, 5, and 8. Genotypes with different letters represent the percentage of stomata with one and two abnormal subsidiary cells that are significantly different from each other (p≤0.05). Statistical analysis is performed based on the Tukey Kramer t-test. All the representative pictures are from leaf 5. Images A, C, E & G are at the same scale. Images B, D, F, & H are at the same scale. Scale bars in G and H are both 50 μm.

The cell divisions that specify the two subsidiary cells in a stomatal complex are independent of each other, therefore any given stomatal complex may have 0 abnormal (i.e., normal), 1 abnormal, or 2 abnormal subsidiary cells. We classified the frequency of abnormal complexes in the 4 genotypes in leaf 3, leaf 5 and leaf 8. (Figure 1). The percentage of stomatal complexes with improper subsidiary cell divisions (and therefore misspecified subsidiary cells) decreased as the leaf number increased. In leaf 3 of *pan1-ems* and *pan2-2*, ∼25% of stomata have abnormal subsidiary cells, but this decreases to ∼20 % in leaf 5 and ∼10% in leaf 8. In leaf 3 of the *pan1;pan2* double mutant, more than 60% of the stomatal complexes have one or two abnormal subsidiary cells, and the percentage of abnormal cells decreases in leaf 5 and 8. This broad range of defective subsidiary cell percentages across different genotypes and leaf numbers, coupled with normal guard cells, provides a useful tool to specifically assay subsidiary cell function during stomatal closure.

To visualize guard cell and subsidiary cell morphology, we generated three-dimensional images of the stomata in four genotypes using confocal microscopy. In *pan1-ems*, *pan2-2*, and *pan1;pan2*, there are always two guard cells that have divided correctly, and the guard cells have a characteristic dumbbell shape. In both wild type and the mutants, the central rod has a thick lignified cell wall, and the guard cell ends are bulbous and have thinner walls (Figure 1). As previously described, *pan2* (but not *pan1*) mutants have “flat side” phenotype, where the outer tips of the subsidiary cells are less pointed (Sutimantanapi *et al*., 2014). Previously, it was shown that guard cells are shorter in the *B. distachyon* allele of *pan1* (Zhang *et al*., 2022). We quantified guard cell length and found it to be shorter in all three mutants using confocal microscopy images (Supplemental Figure 1). Importantly, *pan1, pan2* and *pan1;pan2* all had similarly short guard cell lengths. Moreover, guard cells from stomatal complexes where both subsidiary cells divided normally also had shorter guard cells. This implies that the shorter guard cell length is likely not (solely) due to the effect of a neighboring subsidiary cell that is misspecified, but rather could be due to indirect effects. An important control in our experimental design is the comparison of multiple genotypes, and stomata with 0, 1, or 2 abnormal cells.

### Loss of PAN2, but not PAN1, impairs stomatal closure as inferred by gas exchange

We hypothesized that if subsidiary cells play a critical role in maize stomatal function, the *pan1* and *pan2* single mutants would have moderate defects in stomatal closure. We predicted that the *pan1;pan2* double mutant would exhibit more severe defects in stomatal closure, since *pan1;pan2* has more defective stomatal complexes. Furthermore, we predicted that the earliest emerging leaves, which have the highest frequency of abnormal subsidiary cells, would have the strongest stomatal closure defect; and that later emerging leaves would have subtle or no stomatal closure defects. We assayed stomatal closure by measuring *g_s_* via gas exchange measurements in wild type and mutant plants in response to the darkness (Figure 2A-C; Supplemental Figure S2). We measured *g_s_* in leaf 3, leaf 5 and leaf 8 in B73 (wild type control), *pan1*, *pan2*, and *pan1;pan2.* Graphs with normalized g_s_ (relative to g_s_ at time=0) are shown in Figure 2, and raw (non-normalized) values of g_s_ are shown in Supplemental Figure S2. All data values are listed Supplemental Tables S2 and S3. In B73 control plants, *g_s_* shows a characteristic inverted sigmoidal curve that indicates stomatal closure in response to darkness. As predicted, *pan1;pan2* double mutants showed a stomatal closure defect in response to darkness across all leaves (Figure 2). Moreover, *pan1;pan2* showed milder defects as the leaf number increased, also consistent with our hypothesis. Surprisingly, *pan1* and *pan2* showed different responses from each other, even though they have a similar percentage of abnormal subsidiary cells. Consistent with our prediction, *pan2* showed a defect in stomatal closure that is milder than *pan1;pan2*. On the other hand, *pan1* had no defect in stomatal closure. Importantly, even though *pan1* has no observable defect in dark-induced stomatal closure on its own, the *pan1;pan2* double mutant has a more severe impairment than *pan2-2.* This indicates that even though *pan1* did not have a detectable stomatal closure phenotype as measured by *g_s_*, subsidiary cell specification as influenced by *pan1* is still important for stomatal closure.

**Figure 2.**
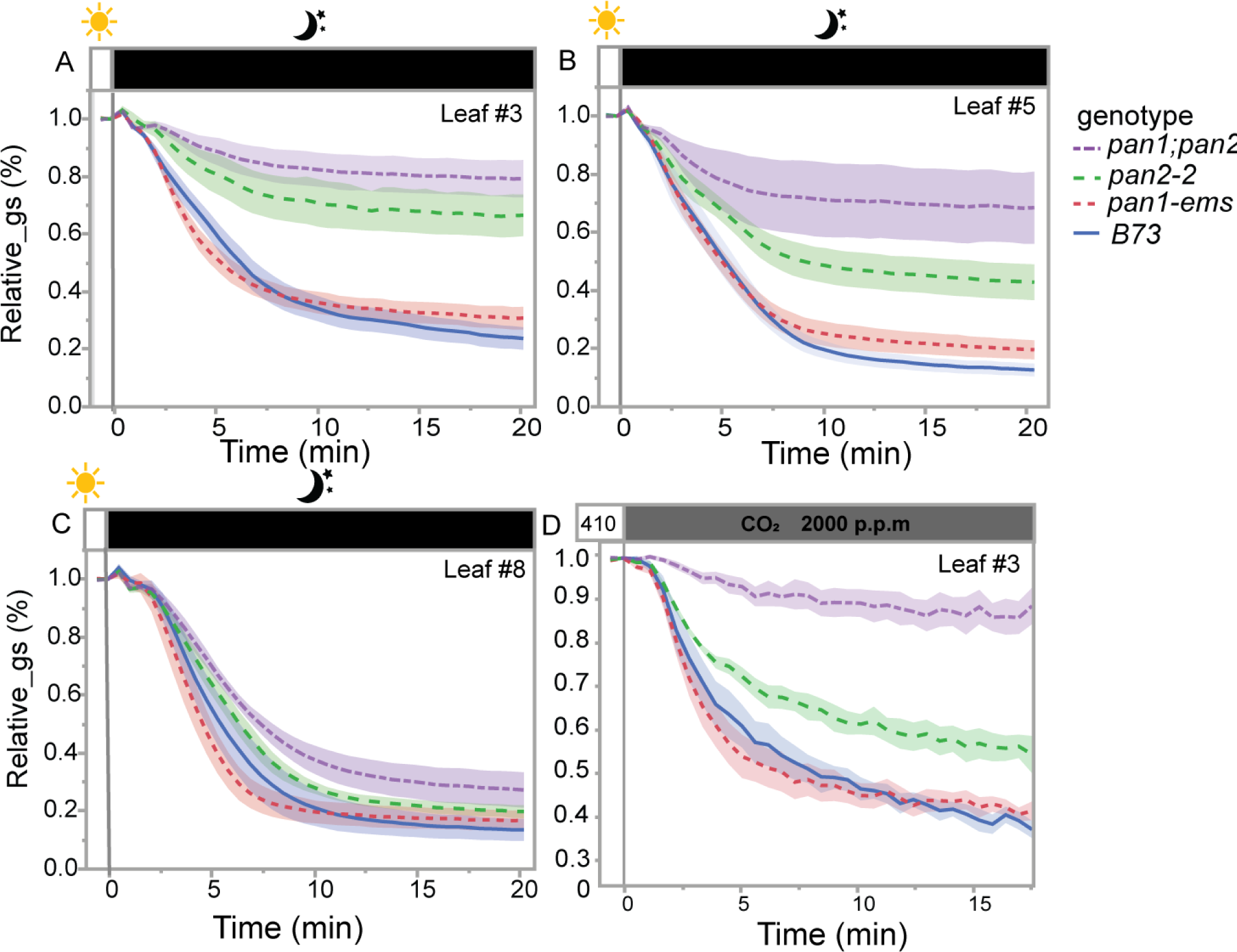
Stomatal closure is impaired in *pan2* and *pan1;pan2,* but is normal in *pan1*. Normalized stomatal conductance (*g_s_*) of B73, *pan1*, *pan2*, and *pan1;pan2* in response to darkness in leaf 3 (A), leaf 5, (B), or leaf 8 (C) was measured over time. PPFD was changed from 1000 to 0 at time zero. (D) Normalized stomatal conductance (*g_s_*) in leaf 3 in response to elevated CO_2_ was measured over time. The CO_2_ concentration of from 410 ppm to 2000 ppm at zero time. Measurements were taken every 5 seconds. Relative *g_s_* was computed for each measured plant by normalizing *g_s_* to the observed steady initial *g_s_* value. The data presented are the mean ± SE of at least 5 individual plants.

To determine if the stomatal closure defect in *pan2* and *pan1;pan2* was specific to darkness-induced closure or a more general defect, we tested another environmental cue that promotes stomatal closure. If the defect is caused by defective subsidiary cells, we reasoned the defect should persist independent of the cue. We increased CO_2_ concentration from 410 ppm (parts per million) to 2,000 ppm to induce stomatal closure. Similar to stomatal closure induced by darkness, *pan2* and *pan1;pan2* showed moderate and strong defects in stomatal closure, respectively, in response to elevated CO_2_ (Figure 2). We did not observe a defect in stomatal closure in *pan1* in response to darkness or elevated CO_2_ (Figure 2).

We confirmed the defect in stomatal closure in *pan2* by analyzing a second, weaker allele of *pan2* (Supplemental Figure S3). The *pan2-O* mutant contains a missense mutation that results in a Ser to Phe change near the N terminus of the protein, which results in less PAN2 protein (Zhang et al., 2012). The percentage of abnormal subsidiary cells in *pan2-O* is less than in *pan2-2* (Supplemental Figure S3). Both alleles showed impaired stomatal closure relative to B73, and *pan2-O* shows a milder defect than *pan2-2* (Supplemental Figure S3. These results are consistent with *pan2-O* being a weaker allele (Zhang et al., 2012).

We favored relative g_s_ (rather than raw values of g_s_) to assay closure, as steady-state g_s_ is influenced by seasonal variation, and we grew our plants in greenhouses with supplemental lighting (and not growth chambers). In *B. distachyon*, seasonal variation has effects on grass stomatal anatomy and morphology, which in turn can have varying effects on assimilation and g_s_ (Nunes *et al*., 2022). We always grew mutant and wildtype plants side-by-side, allowing for comparisons of steady state gs, with the caveat that there is likely environmental influence. In leaf 3, initial g_s_ was higher in *pan2* and *pan1;pan2*, in both our dark-induced closure and CO_2_-induced closure experiment (Supplemental Figure 3). This higher g_s_ that we observed in *pan2* and *pan1;pan2* reduced or disappeared in later emerging leaves. On the other hand, initial g_s_ was lower in *pan1* than in wild type. Similarly, lower steady state g_s_ has been shown for the *B. distachyon* allele of *bdpan1* and *bdpolar* (Zhang *et al*., 2022), which was correlated with a decreased guard cell length in *bdpan1* and *bdpolar* (Zhang *et al*., 2022). We similarly see shorter guard cells in *pan1, pan2* and *pan1;pan2*, and yet only *pan1* shows an initial lower g_s_ (Supplemental Figure 2; Supplemental Figure 3). To ensure the underlying causes were not due to differences in stomatal density, we measured density in the same plants we used to perform the gas exchange assays (Supplemental Figure S4). Stomatal density did not differ in leaf 3 or 8 amongst any genotypes, and if anything was very slightly higher in leaf 5 in *pan1*. The differences in initial g_s_ and stomatal closure that we observe between *pan1* and *pan2* cannot be attributed to size or density. These observations suggest that *pan1* and *pan2* may have underlying fundamental differences. To try and understand how *pan1* and *pan2* may differ, we carefully examined stomatal responses of individual stomatal complexes and cells.

### PAN2 contributes to stomatal function in two distinct ways

Our observation that *pan2*, but not *pan1*, has an effect on stomatal closure was perplexing, given they have a similar percentage of defective stomata. Stomatal gas exchange (*g_s_*) is assayed on a population scale (many thousands of stomatal complexes). Notably, leaf 8 has almost normal stomatal closure, even in the double mutant. It is plausible that on a population level, impairment of stomatal closure of individual defective stomata are masked when the proportion of abnormal stomata is low, due to compensatory responses of the plant. To determine if individual stomata with 0, 1, or 2 abnormal subsidiary cells are able to close normally, we treated leaf segments from mature leaf 5 with ABA and assessed the impact of subsidiary cells on stomatal aperture by confocal microscopy (Figure 3). We measured stomatal aperture in leaf 5 (Figure 3) and leaf 8 (Supplemental Figure S5). Because of the substantial number of comparisons, we included summary data statistics in Supplemental Table S4 and results of pairwise non-parametric tests in Supplemental Table S5. Bonferroni-corrected p-adj values are shown for the +/- ABA comparisons. Wild type stomata in B73 decreased in aperture upon treatment with ABA in both leaf 5 and 8 (Figure 3; Supplemental Figure S5). Consistent with our observation, normally formed stomata in *pan1-ems* also responded to ABA in both leaf 5 and leaf 8. However, in *pan1* mutants with 1 or 2 abnormal subsidiary cells, the aperture prior to ABA treatment was slightly larger, and there was no appreciable decrease in stomatal aperture upon ABA treatment (Figure 3; Supplemental Figure S5); Supplemental Tables S4 and S5). This indicates that even though *pan1* did not show a phenotype on a population level, individual stomata with defective subsidiary cells fail to close properly, and supports the hypothesis that subsidiary cells directly participate in stomatal closure.

**Figure 3.**
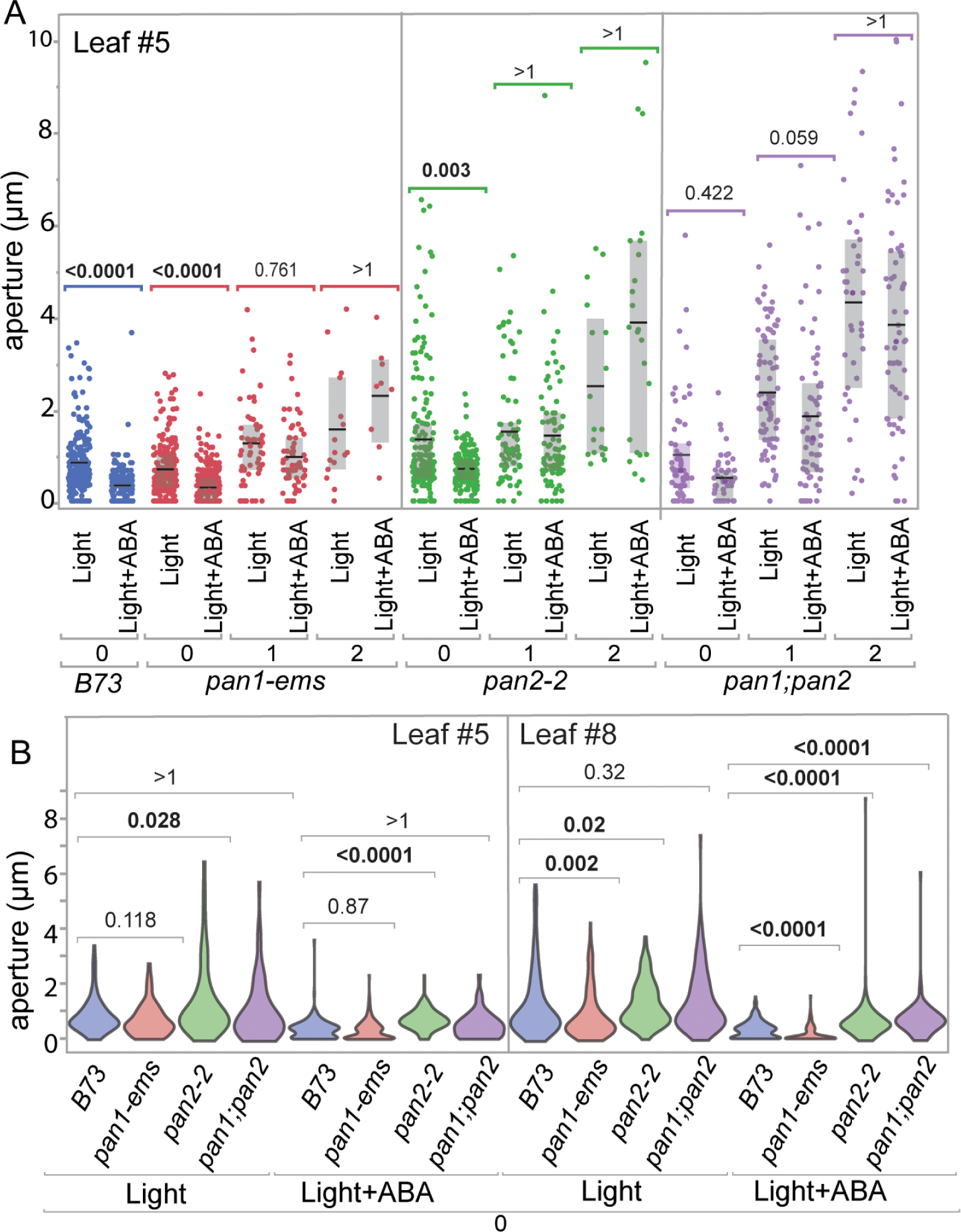
Stomatal aperture of different genotypes in response to ABA. Fully expanded leaf 5 or leaf 8 segments from B73, *pan1, pan2,* or *pan1;pan2* were incubated in opening buffer under 500 μm/m_2_/s light, without or with the addition of 1 mM ABA. (A) Stomata from leaf 5 were classified as having 0 abnormal (i.e., normal), 1 abnormal, or 2 abnormal subsidiary cells, and stomatal apertures were measured. The same experiment using leaf 8 is shown in Supplemental Figure S5A. Each data point represents a single stomatal complex, with 3-5 plants per genotype & treatment used (B) Violin plots showing a subset of the same data from panel A, representing leaf 5 (left panel) and leaf 8 (right panel), is plotted to compare the apertures from stomata with 0 abnormal subsidiary cells. Bonferroni-corrected P-values (P_adj_) values from a from pairwise Wilcoxon signed rank test are shown. Summary statistics are provided in Supplemental Table S2. All pairwise confidence limits, P-values, and adjusted P-values are provided in Supplemental Table S3.

In *pan2* and *pan1;pan2* mutants, stomata with 1 or 2 aberrant subsidiary cells fail to respond to ABA, similar to *pan1.* This is true for both leaf 5 (Figure 3) and leaf 8 (Supplemental Figure S5). Additionally, in *pan2* mutants, stomata with 0 abnormal subsidiary cells respond normally to ABA in both leaves. In leaf 5 of *pan1;pan2* mutants, stomata with 0 abnormal subsidiary cells do not appear to respond (P_adj_ is 0.42; before the Bonferroni correction p=0.02); in leaf 8 *pan1;pan2* stomata with 0 abnormal cells respond robustly while stomata with abnormal cells do not (Supplemental Figure S5).

Notably, *pan2* mutants show a significantly larger aperture in both the absence or presence of ABA relative to both *pan1* and *B73* (Figure 3). To compare normally divided stomata across the 4 genotypes more easily, we replotted a subset of the data from leaf 5, and also included leaf 8 data, using violin plots (Figure 3B). In stomata with normal subsidiary cells (0 abnormal cells), aperture of *pan1* was smaller, in the presence or absence of ABA. This potentially explains why *pan1* mutants have a lower steady state g_s_. On the other hand, *pan2* and *pan1;pan2* stomata had a larger aperture in stomata with normally divided subsidiary cells. Supplemental Figure S5B shows a similar pattern, when stomata with 0, 1, or 2 abnormal subsidiary cells are plotted. This indicates that PAN2 has an additional role in regulating stomatal aperture, independent of the roles PAN1 and PAN2 play during formative subsidiary cell divisions. Together, this indicates PAN2 regulates grass stomatal closure in two ways. Firstly, PAN1 and PAN2 both promote correct subsidiary cell formation during development, which is essential for stomatal closure. Secondly, PAN2 has an additional function in regulating stomatal aperture in all stomata, regardless of whether the subsidiary cell has divided correctly, via an unknown mechanism.

### Subsidiary cells are essential for stomatal closure

Our stomatal aperture measurements in response to ABA allowed us to directly observe stomata with normal and abnormal subsidiary cells. We noticed that in stomata with one abnormal subsidiary cell and one normal cell, the guard cells tended to be curved away from the normal subsidiary cell (Figure 4; Supplemental Figure S6). This was true in all three mutants. In *pan1;pan2*, we observed that 75% (76/101) of guard cells are curved in stomatal complexes with one abnormal subsidiary cell; and all of the curved subsidiary cells are away from the correct subsidiary cells (Figure 4). Similarly, 68% (13/19) of *pan1* and 73% (41/65) of *pan2* stomatal complexes with a single defective subsidiary cell curved away from the morphologically normal cell. Supplemental Figure S6 highlights an example from *pan2-2*, where two stomatal complexes share a single misspecified cell. The two stomata curve in opposite directions; in both cases away from the normally divided subsidiary cell. This qualitative observation implies a pushing or positive force is generated from the normally formed subsidiary cell that results in stomatal pore closure, and suggests a direct mechanical role for subsidiary cell function.

**Figure 4.**
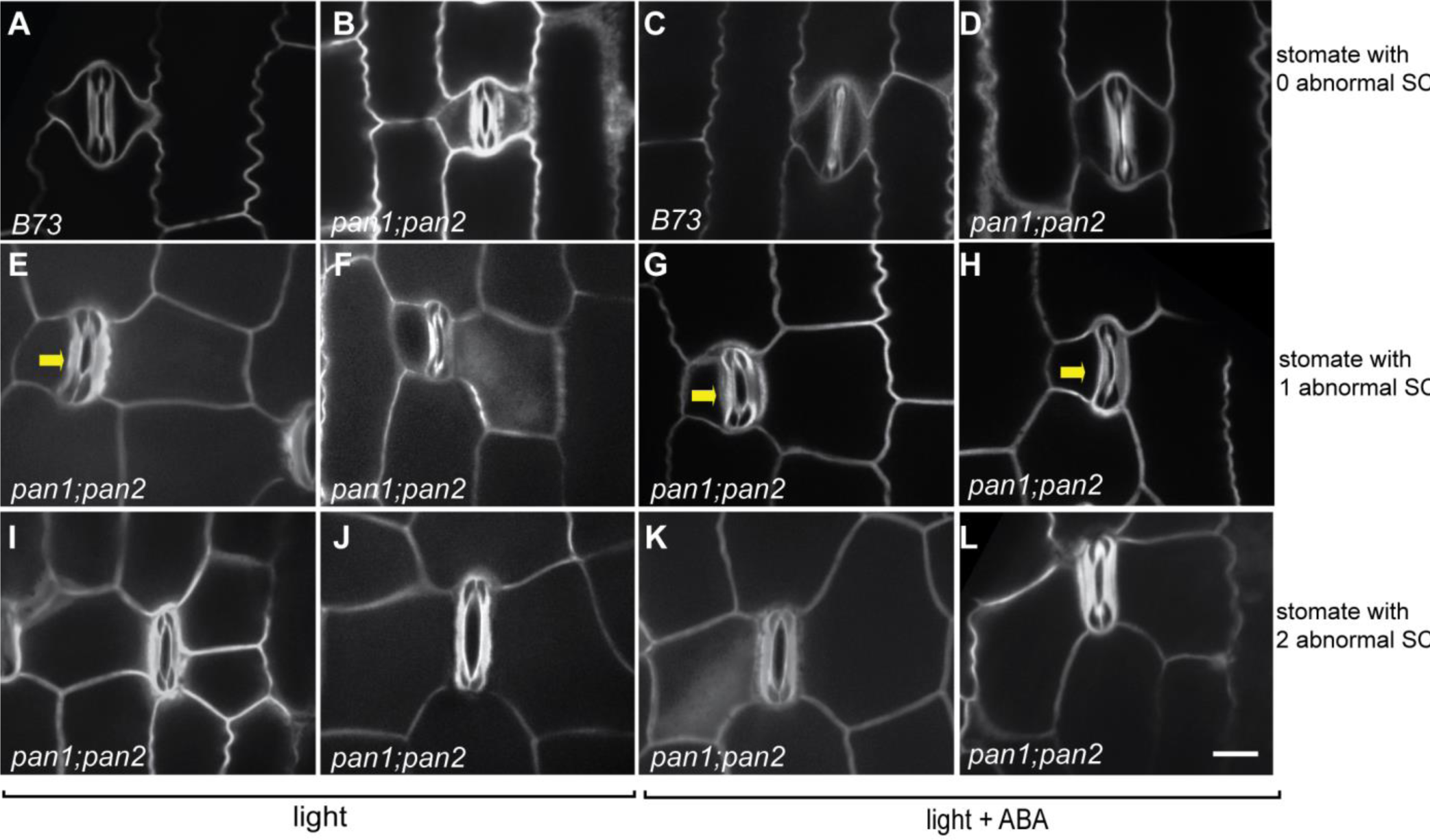
Curved guard cells imply that subsidiary cells exert a positive force. Leaf pieces of fully expanded leaves were treated in opening buffer in the presence or absence of 1 mM ABA. The stomata are classified into three categories based on the subsidiary cell morphologies. Stomata with 1 or 2 abnormal subsidiary cells failed to close in response to ABA treatments. The guard cells in the stoma complex with 1 abnormal subsidiary cell are curved, and all the curves are away from the normally formed subsidiary. Additional images are shown in supplemental Figure S8. All the images are at the same scale; the scale bar in L represents 20 μm.

### PAN2, but not PAN1, is expressed in mature stomata

We further investigated the role of PAN2 in stomatal closure. The one-to-one orthologue of PAN2 in *A. thaliana* is *GUARD CELL HYDROGEN PEROXIDE RESISTANT*

*1 (GHR1)* (Hua *et al*., 2012; Sierla *et al*., 2018; Man *et al*., 2020). In mature leaves, *GHR1* is expressed in guard cells but not in other mature epidermal cells (Hua *et al*., 2012). GHR1 regulates stomatal closure in response to several environmental cues including hydrogen peroxide, light, and ABA (Hua *et al*., 2012). GHR1 might activate the S-type anion channel SLAC1 (Hua *et al*., 2012; Sierla *et al*., 2018). There is no evidence indicating that GHR1 plays a role in stomatal development. Our observation that PAN2 (but not PAN1) is required for stomatal closure, even in correctly divided subsidiary cells, suggests that PAN2 plays a functional role in mature stomata. One possible explanation is that the “flat sides” of *pan2* stomata interfere with the mechanics of closure. An alternative is that PAN2 has a function similar to GHR1, and is present and functionally relevant in mature stomata. Is PAN2 expressed in mature stomata like GHR1? If so, is PAN2 present in guard cells or subsidiary cells? We compared PAN1 and PAN2 expression patterns in developing and mature stomata.

We examined leaves expressing PAN1-YFP and PAN2-YFP at multiple developmental stages using previously characterized marker lines (Humphries *et al*., 2011; Zhang *et al*., 2012; Sutimantanapi *et al*., 2014). PAN1-YFP and PAN2-YFP expression patterns were characterized during stomatal divisions and cell expansion, but their expression in mature stomata was not (Cartwright *et al*., 2009; Zhang *et al*., 2012; Sutimantanapi *et al*., 2014). At the base of a partially expanded immature maize leaf, formative divisions that create stomata occur (Facette *et al*., 2015). There is a developmental gradient from the base of the leaf to the tip, with fully functional stomata at the leaf tip. We examined PAN1-YFP and PAN2-YFP expression along the leaf base at different points. Consistent with past findings, PAN1-YFP and PAN2-YFP polarize in pre-mitotic subsidiary mother cells at the site of guard mother cell contact, and remain polarized after division (Figure 5; Cartwright *et al*., 2009; Zhang *et al*., 2012; Sutimantanapi *et al*., 2014). During cell differentiation and expansion, both PAN1-YFP and PAN2-YFP are expressed, again consistent with previous results (Sutimantanapi *et al*., 2014). PAN1-YFP was undetectable in the leaf tip and mature leaves, with the only signal coming from autofluorescence (Figure 5). Note the heavily lignified guard cells autofluoresce strongly (Supplemental Figure S7). At the tip of partially expanded leaf 5 and in fully expanded leaf 5, PAN2-YFP was clearly expressed in subsidiary cells and pavement cells (Figure 5). PAN2-YFP is no longer polarized in subsidiary cells, but is still present. Due to the guard cell autofluorescence, it was difficult to definitively conclude whether PAN2-YFP was present in guard cells. To address this, we performed plasmolysis experiments. Again, PAN2-YFP was clearly expressed in subsidiary cells, and not seen in the central portion of the guard cells (the central rod of the barbell) (Supplemental Figure S7). We sometimes saw fluorescence in the bulbous ends of the guard cells, but this was inconsistent and difficult to resolve from autofluorescence. Therefore, we concluded that like GHR1, PAN2 is expressed in the stomatal complex. PAN2 is expressed in subsidiary cells, and may also be expressed in guard cells. Our PAN1-YFP data indicates that PAN1 is not expressed in mature stomatal complexes (Figure 5).

**Figure 5.**
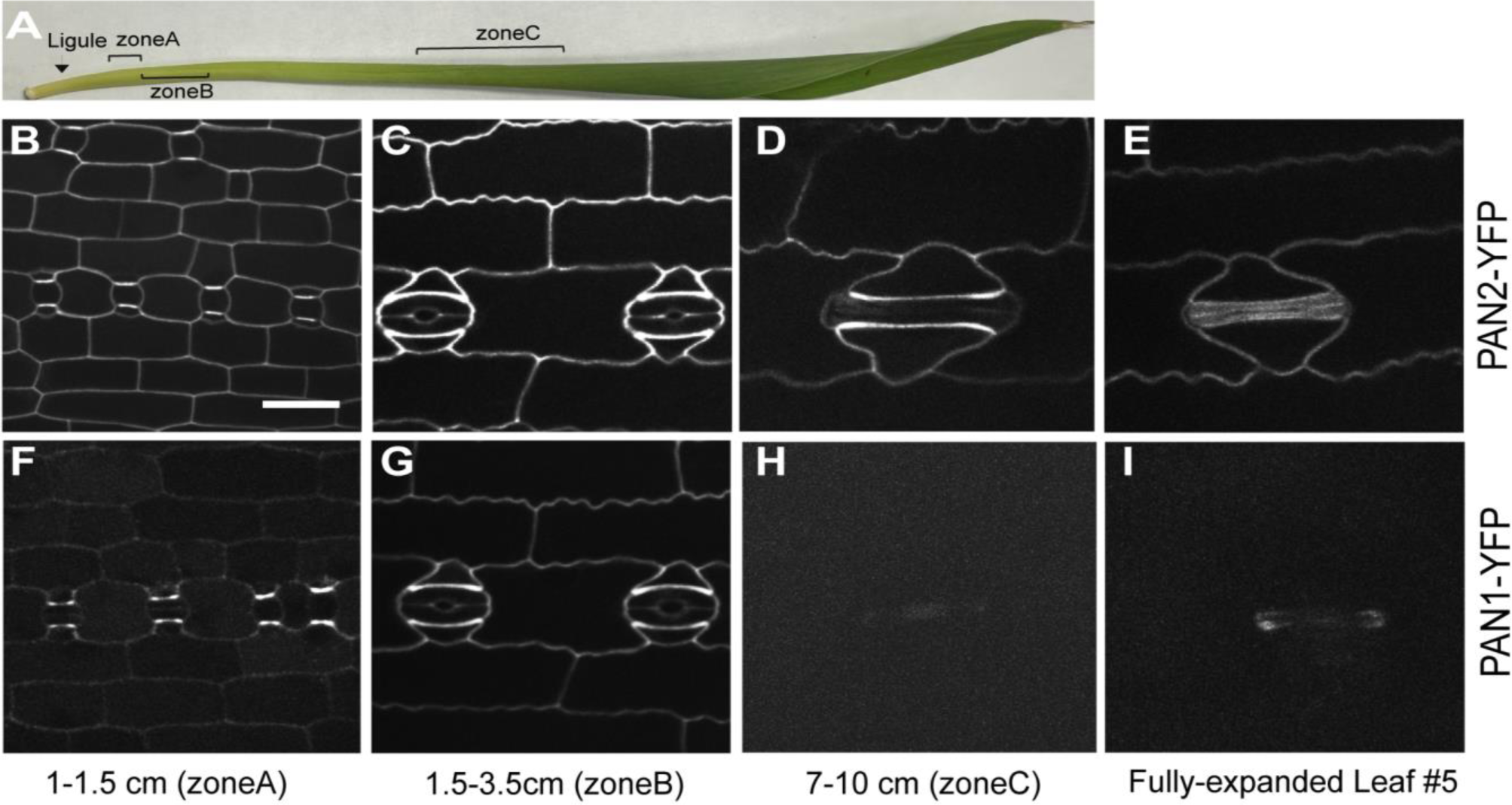
PAN1 and PAN2 have different expression patterns in division zones and fully expanded leaves. The outer leaves of 15-d-old plants expressing either PAN2-YFP (B-D) or PAN1-YFP (F-H) were removed to expose partially expanded leaf 5, where the developing ligule was within 0.5 cm of the leaf base. Leaf 5 was cut into 3 sections as illustrated in A to examine expression in partially expanded leaves. The leaf blade of a fully expanded leaf 5, where the leaf sheath and leaf blade are no longer growing, was examined for PAN2-YFP (E) or PAN1-YFP (I) expression. All images are at the same scale; the scale bar in A represents 20 μm.

We further corroborated the expression of PAN2 and PAN1 in leaves via qPCR. Previous proteomic data indicated the presence of both proteins in immature leaves, but neither was detected in mature leaves (Facette *et al*., 2013). This could be due to a low abundance of PAN2 in mature leaves, coupled with the relative insensitivity of proteomic detection. Similarly, neither *Pan1* nor *Pan2* were detected in single nuclear RNA-sequencing (snRNA-seq) experiments with mature maize leaves (Sun *et al*., 2022); however *Pan2* was also not detected via snRNA-seq in the stomatal division zone where it is known to be expressed (Zhang *et al*., 2012; Sun *et al*., 2022). Genes expressed at low levels are also often below the level of detection in single cell/single nucleus experiments. We performed quantitative real-time PCR (qRT-PCR) in different leaf numbers and division zones, using the mutants as a negative control. Both *PAN1* and *PAN2* are expressed in leaves where stomatal divisions and cell expansion are occurring (Supplemental Figure S8; Supplemental Tables S6 and S7). PAN2 is expressed in fully expanded leaf 3, 5 and 8, albeit at low levels. Nonetheless, the amount of expression was greater than what was detected in the null mutant. This is consistent with PAN2 having a role in stomatal function after development has completed. PAN1 expression in fully expanded leaves was similar to the null mutant (Supplemental Figure S8; Supplemental Tables S6 and S7). Together, these data indicate PAN2, but not PAN1, is expressed in mature stomata, consistent with a role for only PAN2 during stomatal function.

### PAN2 may regulate stomatal closure via a Ca^2+^-dependent pathway

In *A. thaliana*, *ghr1* is insensitive to dark-, CO_2_-, and ABA-induced stomatal closure (Hua *et al*., 2012), similar to *pan2*. Application of exogenous Ca^2+^ results in stomatal closure in wild type plants and restores stomatal closure in *ghr1,* which indicates that Ca^2+^ acts downstream of GHR1 in A. *thaliana* (Hua *et al*., 2012; Sierla *et al*., 2018). Given the phenotypically similar impairment of stomatal closure observed in *ghr1* and *pan2*, we wanted to test if the *pan2* phenotype could also be rescued by exogenous application of Ca^2+^. We treated epidermal peels of B73 and *pan2-2* using the opening buffer previously described, in the presence or absence of 10 mM CaCl^2+^ under 500-700 PPFD light. Consistent with our previous observation, *pan2-2* shows larger stomatal apertures than B73 in both conditions (Figure 6). Application of exogenous Ca^2+^ induced stomatal closure in B73. Strikingly, *pan2* did not respond to application of exogenous Ca^2+^ (Figure 6). This suggests that the molecular pathway that PAN2 participates in may be differently wired in maize vs *A. thaliana,* and that in maize Ca^2+^ is upstream, rather than downstream, of PAN2.

**Figure 6.**
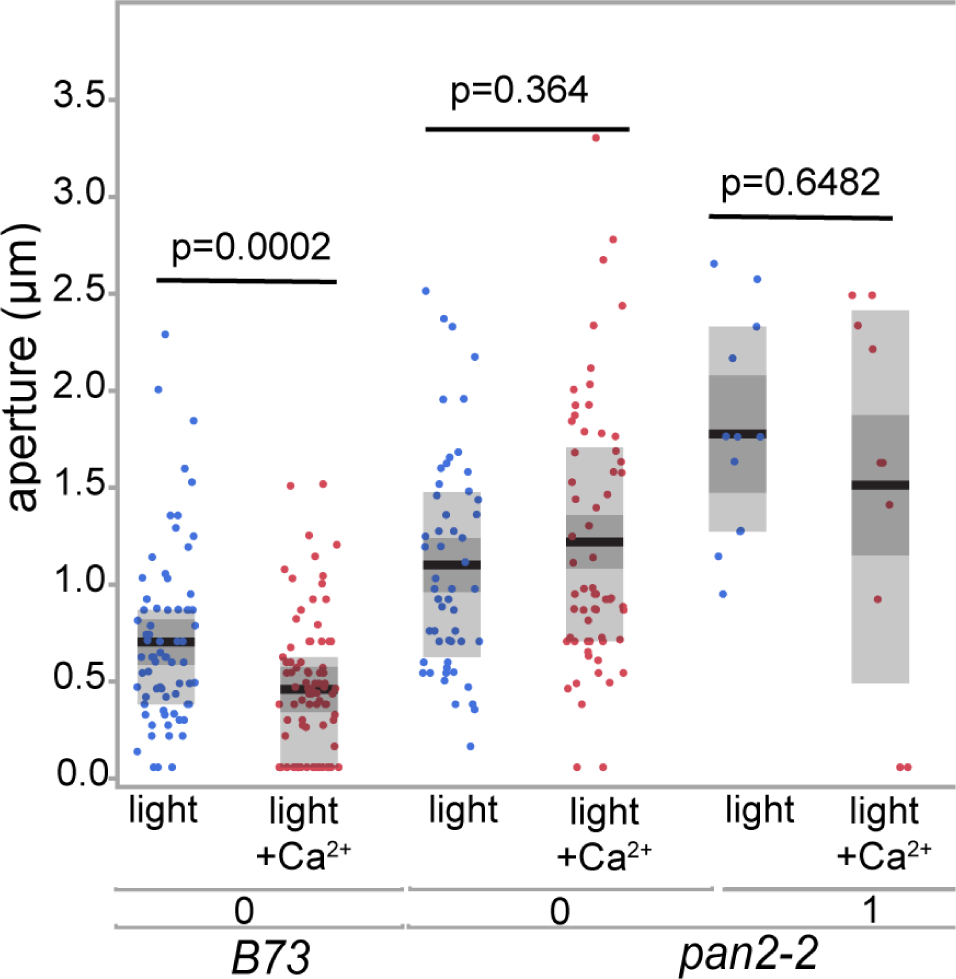
pan2 does not respond to exogenous Ca_2+_. Leaf tissue from *B73* or *pan2* was incubated in opening buffer or buffer supplemented with 10 mM Ca_2+_ under 500 μm/m_2_/s light. After 2 hours, the pieces were stained with propidium iodide (PI) and imaged on the confocal microscope. Stomatal apertures were measured from the images using FIJI. Stomata were classified as having 0 abnormal (i.e., normal) or 1 abnormal subsidiary cell. *p*-values shown are from a Wilcoxon signed rank test.

## Discussion

### Properly formed subsidiary cells are essential for stomatal movements in maize

Stomata facilitate gas exchange for carbon dioxide absorption and drive plant transpiration, but are also the main source of plant water loss. The fast movements of grass stomata are attributed to the combination of subsidiary cells and dumbbell-shaped guard cells (Franks and Farquhar, 2007). Subsidiary cells are not unique to grasses (Gray *et al*., 2020; Nunes *et al*., 2020; Cheng & Raissig, 2023). Substantive evidence across many plant species supports that subsidiary cells, or even neighboring pavement cells, accumulate ions in a manner opposite to guard cells during stomatal movement (Raschke & Fellows, 1971; Willmer & Pallas Jr., 1973; Dayanandan & Kaufman, 1975). However, subsidiary cells do not seem to be necessary for stomatal movements in all species. For example, in *Commelina communi (L.)*, which has lateral and polar subsidiary cells, stomata respond in the absence of living subsidiary cells (Squire & Mansfield, 1972). Franks and Farquhar (2007) convincingly showed that volume changes in grass subsidiary cells oppose the volume changes in guard cells; and there are subsidiary cell-specific ion channels (Majore *et al*., 2002; Büchsenschütz *et al*., 2005). Our gas exchange and microscopic analyses of stomata with abnormally divided subsidiary cells, at varying percentages, confirm that subsidiary cells are essential for stomatal closure in maize. In particular, the striking observation of curved guard cells in stomatal complexes with a single aberrant subsidiary cell indicates a positive force exerted by subsidiary cells that promotes closure. It is important to be aware that *pan2* has a subsidiary cell morphological defect not shared by *pan1* (Sutimantanapi *et al*., 2014). Subsidiary cells in *pan2* have “flat sides”. However, we believe it is less likely this flat-sided defect is the source of the *pan2* stomatal closure defect for two reasons. Firstly, flat-sided subsidiary cells occur naturally in grasses lacking epidermal crenulations, such as *B. distachyon.* Secondly, and more importantly, our microscopic data indicates that in *pan2* and *pan1;pan2* stomatal complexes with a single abnormally divided subsidiary cell, the other subsidiary cell still exerts a positive or pushing force. Future experiments where *pan2* is rescued will cell type specific and developmentally regulated promotors could potentially untangle the effects of PAN2 during subsidiary formative cell divisions vs. subsidiary cell expansion and morphology vs. mature subsidiary cell and/or guard cell function.

Individual abnormal stomata observed microscopically in *pan1* mutants have closure defects, however gas exchange data did not detect a defect. Because *pan1;pan2* double mutants showed a more severe defect than the *pan2* single mutant, *pan1*-induced subsidiary cell misspecification is a factor in stomatal closure. Later emerging leaves in *pan2* and *pan1;pan2,* which have fewer abnormal stomata than early leaves, also show mild to no defect. Hence it is plausible that a certain threshold percentage of abnormal stomata is required to show a population-level defect.

We used changing light (1000 to 0 μmol m^−2^ s^−1^) and changing CO_2_ (410 ppm to 2000 ppm) to induce stomatal closure. In our gas exchange experiments, g_s_ started to plateau shortly after 5 minutes and well within 10 minutes. Even *pan2* and *pan1;pan2* mutants, which do not fully close, plateaued within this time scale. This is comparable to previous values found for maize (McAusland *et al*., 2016), but faster than that found for rice and *B. distachyon* (McAusland *et al*., 2016; Qu *et al*., 2016; Nunes *et al*., 2022; Zhang *et al*., 2022). A meta-analysis of many plants species calculated the time to reach 63% of closure (t_cl_), which was similar (5.9 minutes) across grasses, but longer for non-grass species (>15 minutes) (Vico *et al*., 2011). Several of these other studies used step changes from 1000 to 100 μmol m^−2^ s^−1^, which may be an important variable. The importance of stomatal closure speeds and its relationship to water use efficiency has garnered recent attention – plausibly our data could be a resource for other studies (Lawson & Blatt, 2014; Raven, 2014; Lawson & Vialet-Chabrand, 2019)

### PAN2 is required for stomatal function in maize

Some studies have shown that subsidiary cells may participate in stomatal regulation signaling pathways. The subsidiary-cell-specific expressed glucose transporter CST1 positively regulates stomatal opening (Wang *et al*., 2019b). The *cst1* mutant showed reduced stomatal opening, starvation, and early senescence, suggesting the necessary role for subsidiary cells in stomatal function or signaling mechanisms, although the mechanism behind this is unclear (Wang *et al*.,

2019b). It was also shown that the common signaling messenger H_2_O_2_ accumulated in subsidiary cells and is necessary for drought-induced stomatal closure in maize, demonstrating the significance of stomatal signaling mechanisms (Yao *et al*., 2013; Zhang *et al*., 2020). Once again, the detailed mechanism of the subsidiary cell anatomy, membrane transporters, and their relationship with the guard cell behavior is still unclear. It is clear however that the SLAC1 transporter in grasses functions differently than in *A. thaliana*, as it has a requirement for nitrate (Schäfer *et al*., 2018). How subsidiary anatomy interacts with guard cell anatomy is also surely important. Guard cells in *Bdmute/sid* mutant are shorter, and the dumbbell shape is present but slightly altered (Durney *et al*., 2023). Whether this is because of a non-cell autonomous effect (i.e., a lack of neighboring subsidiary cells) or another effect is unclear; however, both *Bdmute* and *pan* mutants have shorter guard cells. The maize *bzu3* mutant has drastically altered guard cell morphology, which affects stomatal function (Zhou *et al*., 2023).

A strong connection between GHR1 and apoplastic ROS has been suggested (Hua et al., 2012; Devireddy et al., 2018), and genetic analysis indicated that GHR1 acts downstream of ROS production but upstream of Ca^2+^ channel activation (Hua et al., 2012). Maize stomata are similar to *A. thaliana* and other plant species in that exogenous Ca^2+^ and H_2_O_2_ promote stomatal closure (Yao *et al*., 2013; Zhang *et al*., 2020). We showed that *pan2* showed fully impaired stomatal closure in response to supplemental Ca^2+^ suggesting that PAN2 is indispensable for the Ca^2+^-dependent stomatal closure pathway. Co-immunoprecipitation and Bimolecular fluorescence complementation (BiFC) experiments showed that GHR1 physically interacts with AtSLAC1, and perhaps activates SLAC1 (Hua *et al*., 2012; Sierla *et al*., 2018). SLAC1 is a conserved anion channel controlling turgor pressure in guard cells, thereby regulating stomatal movement in response to environmental signals in monocots and dicots (Negi *et al*., 2008; Vahisalu *et al*., 2008). Since GHR1 is a pseudokinase (as is PAN2), its proposed effect on SLAC1 is to act as a scaffold for additional regulators rather than a direct phosphorylation activity (Sierla *et al*., 2018). In both maize and *A. thaliana*, *slac1* mutant plants have impaired stomatal closure (Vahisalu *et al*., 2008; Qi *et al*., 2018). ZmSLAC1 is expressed explicitly in guard cells in maize (Qi *et al*., 2018). Our expression analyses clearly indicated that PAN2 is expressed in subsidiary cells, but PAN2 expression in guard cells was less conclusive. The recent rise in single cell/single nuclear RNA-seq experiments may help resolve PAN2 expression in the stomatal complex. Recently, Sun et al. (2022) single-nucleus RNA sequencing to identify gene expression in guard cells and subsidiary cells. While this technique in general and this paper specifically is a powerful resource a general limitation of single-cell RNA sequencing that often genes with low expression can sometimes be overlooked, and that closely related genes sometimes not be distinguished.

Since guard cells and subsidiary cells have opposing turgor, and *ghr1* and *pan2* both have defects in stomatal closure, it would make sense if PAN2 action was primarily in the guard cell (rather than the subsidiary cell) - assuming the molecular function of GHR1 and PAN2 is conserved. However, this may or may not be true. PAN2 may function in subsidiary cells - perhaps regulating a SLAC1 paralog. Alternatively, PAN2 may regulate other proteins. Recently, PAN2 interactors were identified via co-immunoprecipitation/mass spectrometry in the stomatal division zone (Nan *et al*., 2023). Amongst the high-confidence interactors were several maize proteins similar to *A. thaliana* AHA family proton transporters and a protein kinase in the SnRK family, both of which are important for stomatal function in *A. thaliana* (Nan *et al*., 2023). Perhaps PAN2 (and GHR1?) have more general roles regulating other transporters and proteins. Both PAN2 and GHR1 are expressed in other tissues, including meristems (Hua *et al*., 2012; Zhang *et al*., 2012). An unbiased approach to identifying interactors in both stomata and other tissues will help identify the precise roles of these pseudokinases. Combining such data with future genetic and molecular work will help untangle the similar and dissimilar roles of PAN2 and GHR1. Recent single-cell gene expression analyses of maize stomatal complexes identified subsidiary cell specific genes expressed at different times in development (Sun *et al*., 2022). Future experiments where PAN2 is driven by promotors early in development (during subsidiary cell formation) vs later (in mature stomata) will help untangle the dual roles of PAN2. Coupled with double mutant analyses of guard cell specific and subsidiary cell specific *slac/slah* mutants will help us understand what the similar and dissimilar roles of PAN2 and GHR1 in different species and cell types.

## Acknowledgments

We thank Chris Phillips, Bob Skalbite, Neal Woodward, and other greenhouse and farm staff at the University of Massachusetts Amherst for help in maintaining maize plants. We thank Ahmet Bakirbas and Stavroula Fili from Elsbeth Walker lab for their advice on the qPCR experiments. We thank Laurie Smith for reviewing our work and offering valuable feedback. This material is based upon work supported by the National Institute of Food and Agriculture, U.S. Department of Agriculture, the Center for Agriculture, Food and the Environment and the Biology Department at University of Massachusetts Amherst, under project number MAS00570. The contents are solely the responsibility of the authors and do not necessarily represent the official views of the USDA or NIFA.

## Author contributions

L.L. and M.R.F. conceived the project and designed experiments. L.L., M.A.A., and T.M. performed experiments and analyzed data. L.L. and M.R.F. wrote the manuscript with the input of other authors. M.R.F. supervised the project. All the authors read and approved the final manuscript.

## List of Supplemental Figures and Tables

### Supplemental Figures (available as a PDF)

**Figure S1.** Guard cells in *pan1, pan2,* and *pan1;pan2* are shorter than those in B73 in leaf 5.

**Figure S2.** Stomatal closure is impaired in *pan2* and *pan1;pan2* and but is normal in *pan1* (absolute conductance).

**Figure S3.** Comparison of stomatal developmental defects between *pan2-2* (strong allele) and *pan2-o* (weak allele).

**Figure S4.** Stomatal density in B73, *pan1*, *pan2*, and *pan2;pan1*

**Figure S5.** The stomatal aperture of different genotypes of in response to ABA (leaf 8).

**Figure S6.** Representative images of “curved” guard cells from *pan2-2* and *pan1;pan2* mutants with or without treatment with ABA

**Figure S7.** PAN2 is expressed in subsidiary cells and pavement cells.

**Figure S8:** Relative expression levels of *PAN1* and *PAN2* in division zones and fully expanded leaf 3, 5, and 8.

### Supplemental Tables (available as .xlsx)

**Supplemental Table 1.** Primer sequences for gene expression analysis.

**Supplemental Table S2.** Raw and normalized values of g_s_ and A under different lighting conditions

**Supplemental Table S3.** Raw and normalized values of g_s_ and A under different CO_2_ concentrations.

**Supplemental Table S4.** Summary statistics of stomatal aperture for leaf 5 and leaf 8. **Supplemental Table S5.** Results of pairwise comparisons of statistical tests for stomatal aperture for leaf 5 and leaf 8.

**Supplemental Table S6.** Raw CT values and calculated ratios for qRT-PCR experiments.

**Supplemental Table S7.** Pairwise t-tests of qRT-PCR data.

---

**Fig. S1.**
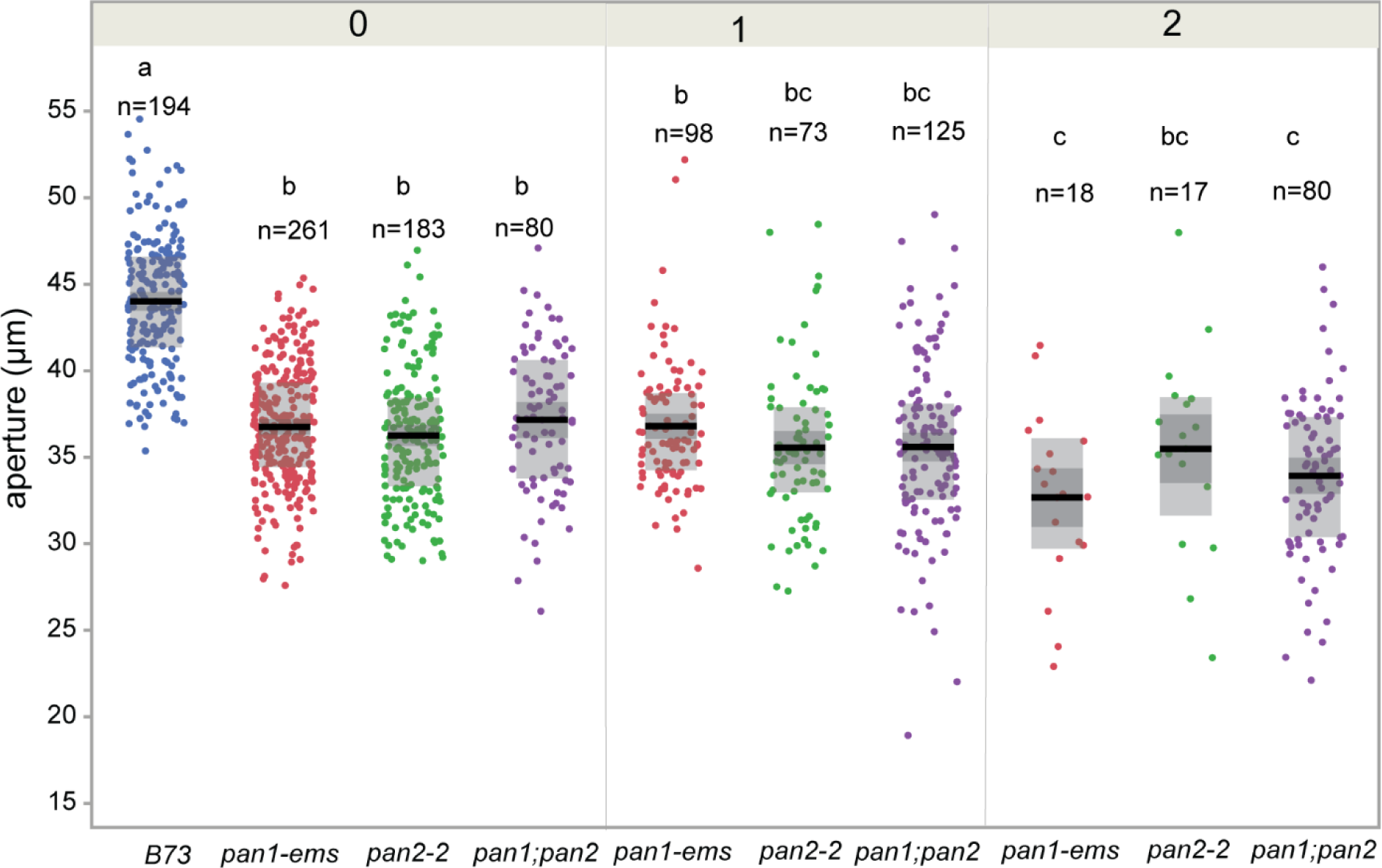
Guard cells in *pan1, pan2,* and *pan1;pan2* are shorter than those in B73, in leaf 5. Guard cell length was measured using confocal images. Stomata were classified as having 0 abnormal (i.e., normal), 1 abnormal, or 2 abnormal subsidiary cells. Each data point represents a single guard cell pairs length, with 3-5 plants per genotype used. n represents the numbers of guard cell pairs used for measurement. Black bars indicate means; dark grey boxes are standard errors and larger light grey boxes indicate interquartile range. The number (0, 1, 2) above the panel represents the number of abnormal subsidiary cells in the stomate. Genotypes with different letters represent the guard cell lengths that are significantly different from each other (p≤0.05) Tukey Kramer Test.

**Fig. S2.**
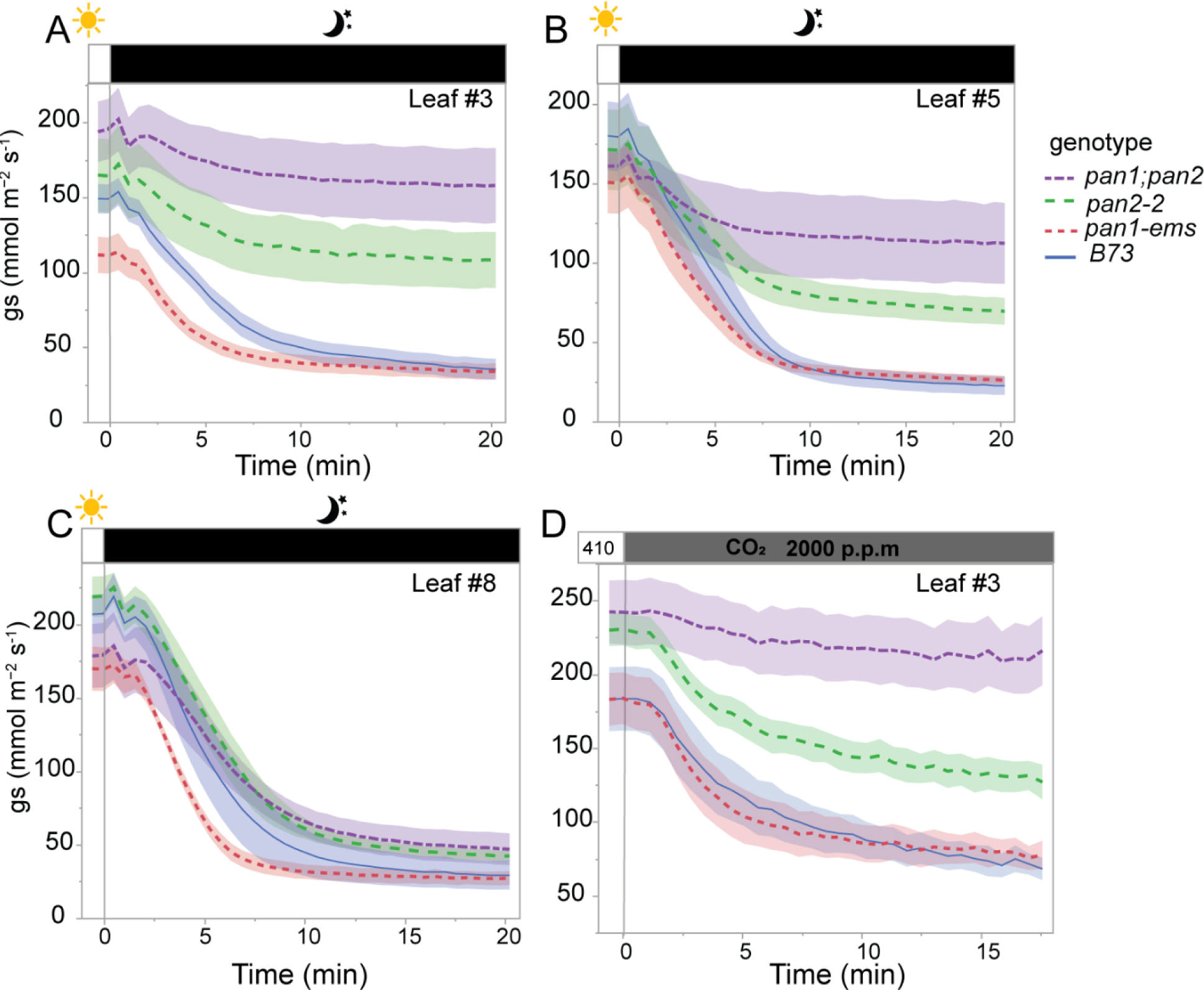
Stomatal closure is impaired in *pan2* and *pan1;pan2* and but is normal in *pan1* (absolute conductance). The normalized stomatal conductance data plotted in Figure 2 are plotted here as absolute stomatal conductance. Absolute stomatal conductance (*g_s_*) of *pan2*, *pan2;pan*1, *pan1*, and B73 in response to darkness in leaf 3 (A), leaf 5, (B), or leaf 8 (C) was measured over time. PPFD was changed from 1000 to 0 at time zero. (D) Normalized stomatal conductance (*g_s_*) in leaf 3 in response to elevated CO_2_ was measured over time. The CO_2_ concentration of from 410 ppm to 2000 ppm at zero time. Measurements were taken every 5 seconds. The data presented are the mean ± SE of at least 5 individual plants. Pertains to Figure 2.

**Fig. S3.**
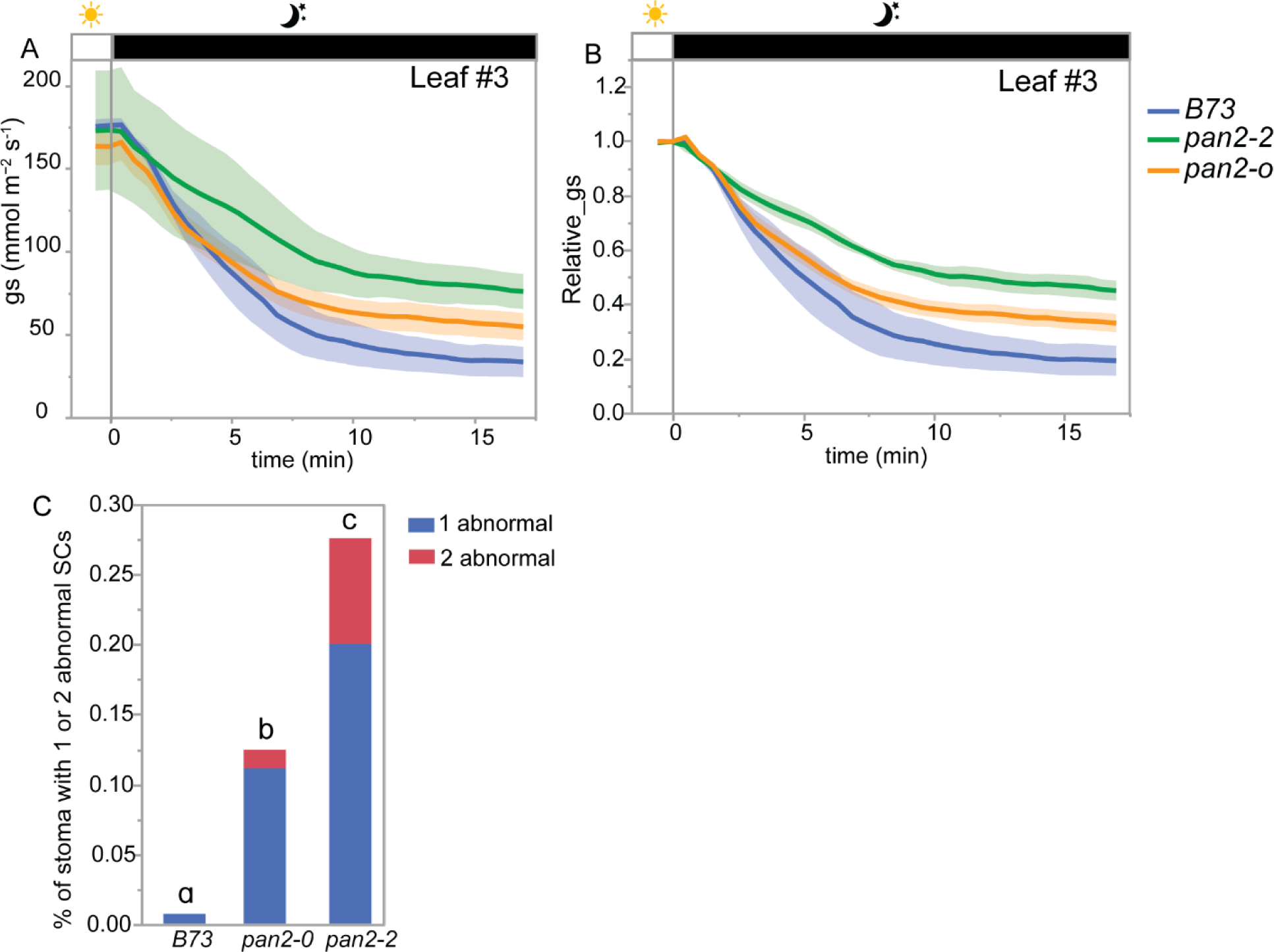
Comparison of stomatal developmental defects between *pan2-2* (strong allele) and *pan2-O* (weak allele). (A) The *g_s_* of B73, *pan2-o*, and *pan2-2 was* measured in leaf 3 using in response to darkness. Light was turned off from 1000 to 0 at zero for the darkness response. The data presented are the mean ± SE of at least 3 individual plants. (B) Relative *g_s_* of B73, *pan2-o* and *pan2-2* computed for each measured plant by normalizing *g_s_* to the observed steady initial *g_s_* value. (C) Percentage of abnormal subsidiary cells (SCs) in B73, *pan2-O*, and *pan2-2* in leaf 3. All the plants were planted simultaneously. At least 300 stomata complexes were counted for each plant, and at least three plants were counted for each genotype. Genotypes with different letters represent the percentage of stomata with one and two abnormal subsidiary cells that are significantly different from each other (p≤0.05). Statistical analysis is performed based on the Student’s t-test. Pertains to Figure 2.

**Fig. S4.**
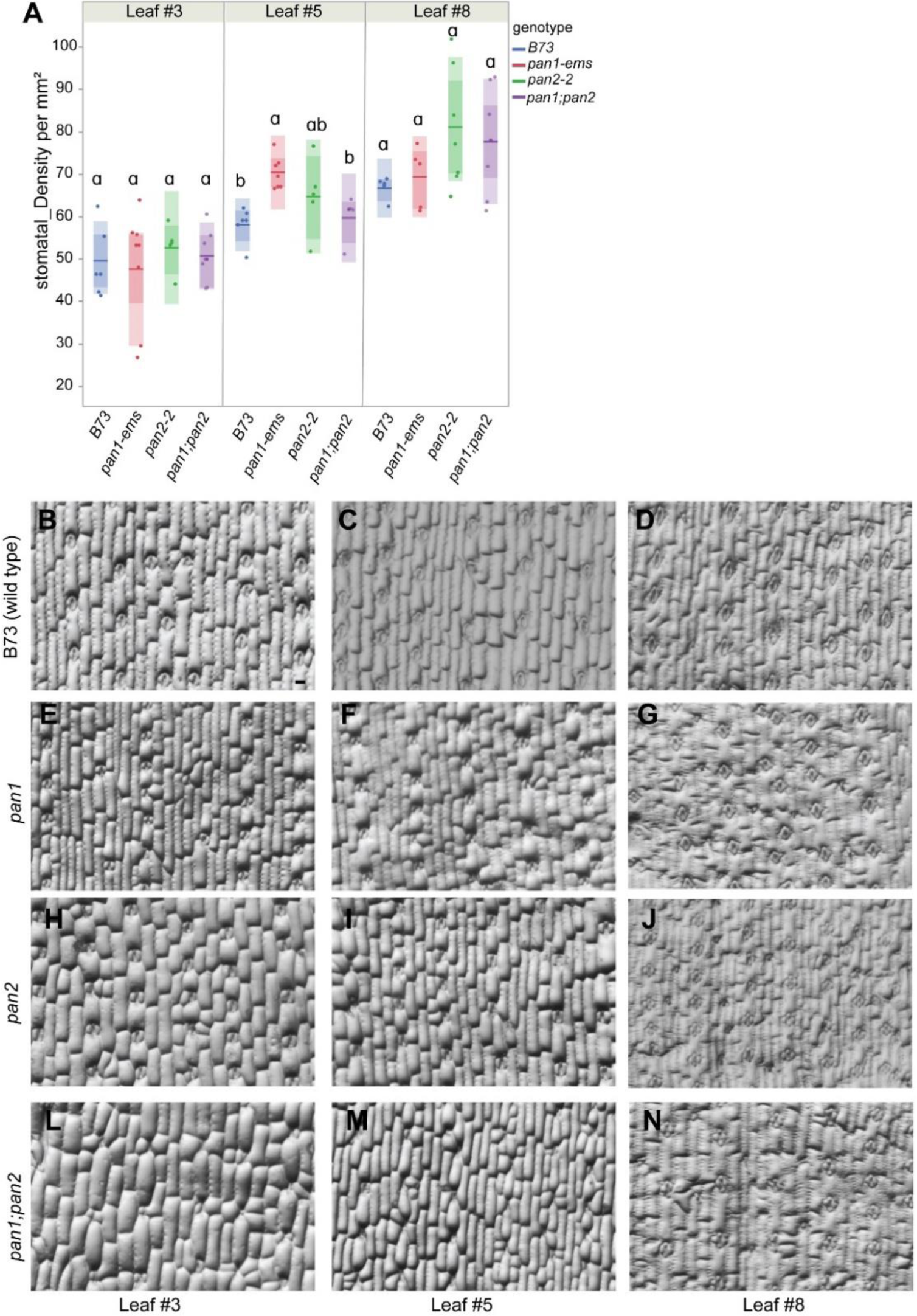
Stomatal density in B73, *pan1*, *pan2*, and *pan2;pan1*. (A) Stomatal density was measured using epidermal glue impressions. Density was measured in the same plants used for conductance measurements. Bars indicate means; dark shaded boxes indicate standard errors and lighter colored boxes indicate interquartile means. Genotypes with different letters indicate densities that are significantly different from each other (p≤0.05) based on Student’s t-test. (B-N) The representative images of each genotype from leaf 3, 5, and 8. All the images were at the same scale. Scale bar in B is 50 μm. Pertains to Figure 2.

**Fig. S5.**
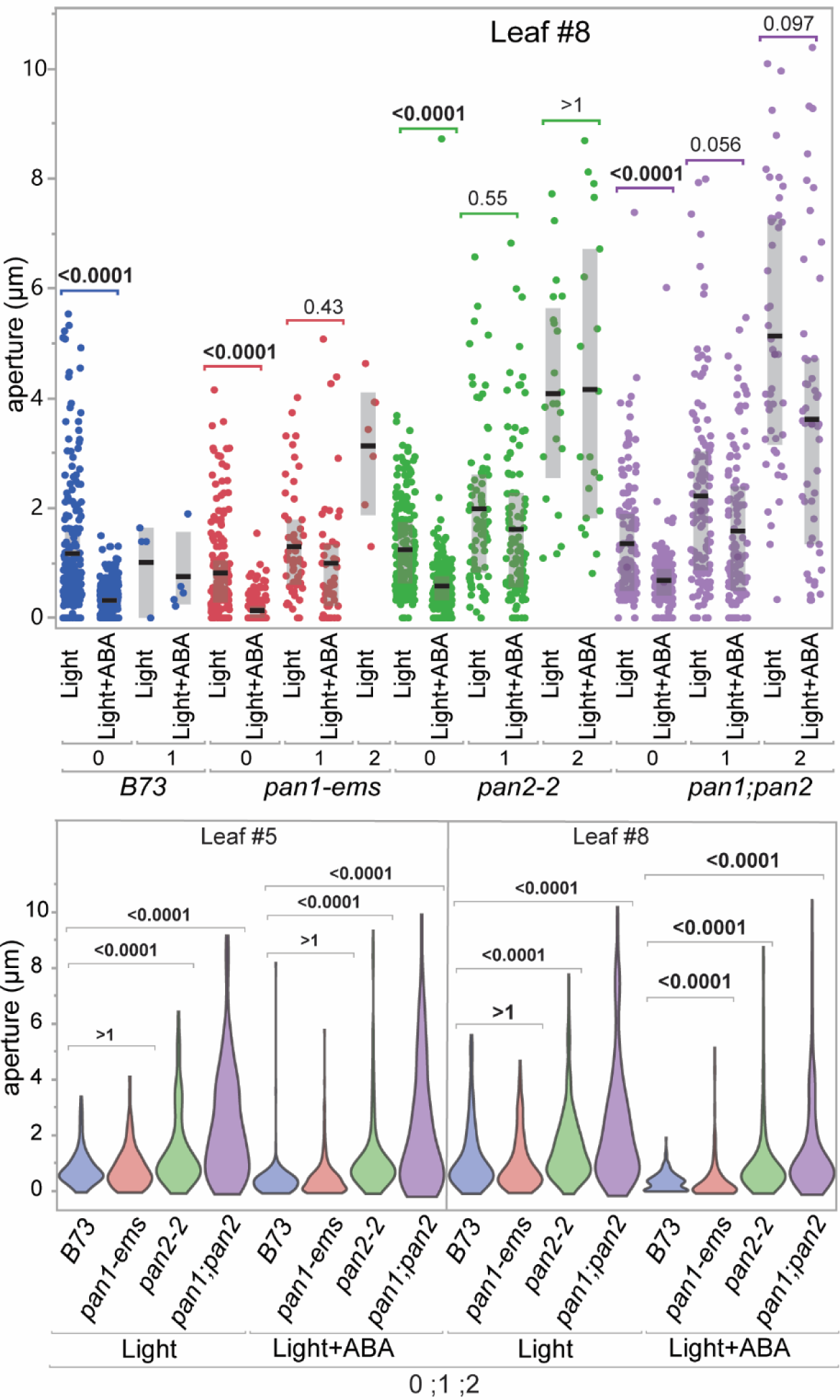
Stomatal aperture of different genotypes in response to ABA. (A) Fully expanded leaf 8 segments from B73, *pan1,* or *pan1;pan2* was incubated in an opening buffer under 500 µm/m^2^/s light, +/- 1 mM ABA. Stomata were classified as having 0, 1, or 2 abnormal subsidiary cells. Each data point represents a single stomatal complex, with 3-5 plants per genotype & treatment were used. Black bars indicate medians; grey boxes indicate interquartile ranges. (B) Violin plots of aperture measurements from all stomates with any number abnormal subsidiary cells, which represents the entire leaf sample (i.e., similar to gas exchange measurements). Leaf 5 is shown in the left panel; leaf 8 is shown on the right. Padj-values from a Bonferroni-corrected Wilcoxon comparison are as labeled. Pertains to Figure 3.

**Fig. S6.**
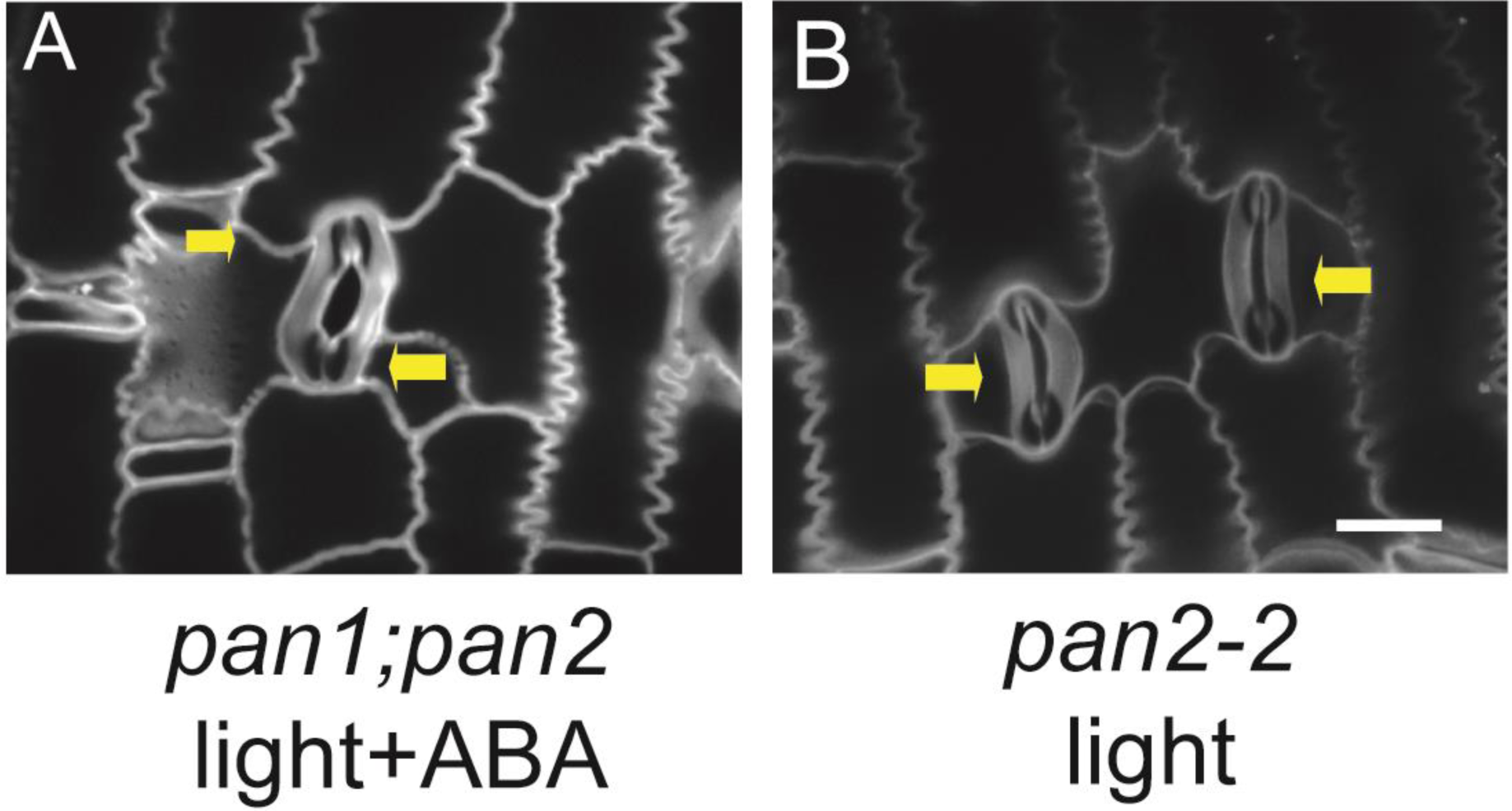
Representative images of “curved” guard cells from *pan2-2* and *pan1;pan2* mutants with or without treatment with ABA. Leaf pieces of fully expanded leaves were treated in opening buffer in the presence or absence of 1 mM ABA. The yellow arrows indicate the direction of the subsidiary cell “push”. The scale bar is 20 μm. Pertains to Figure 4.

**Fig. S7.**
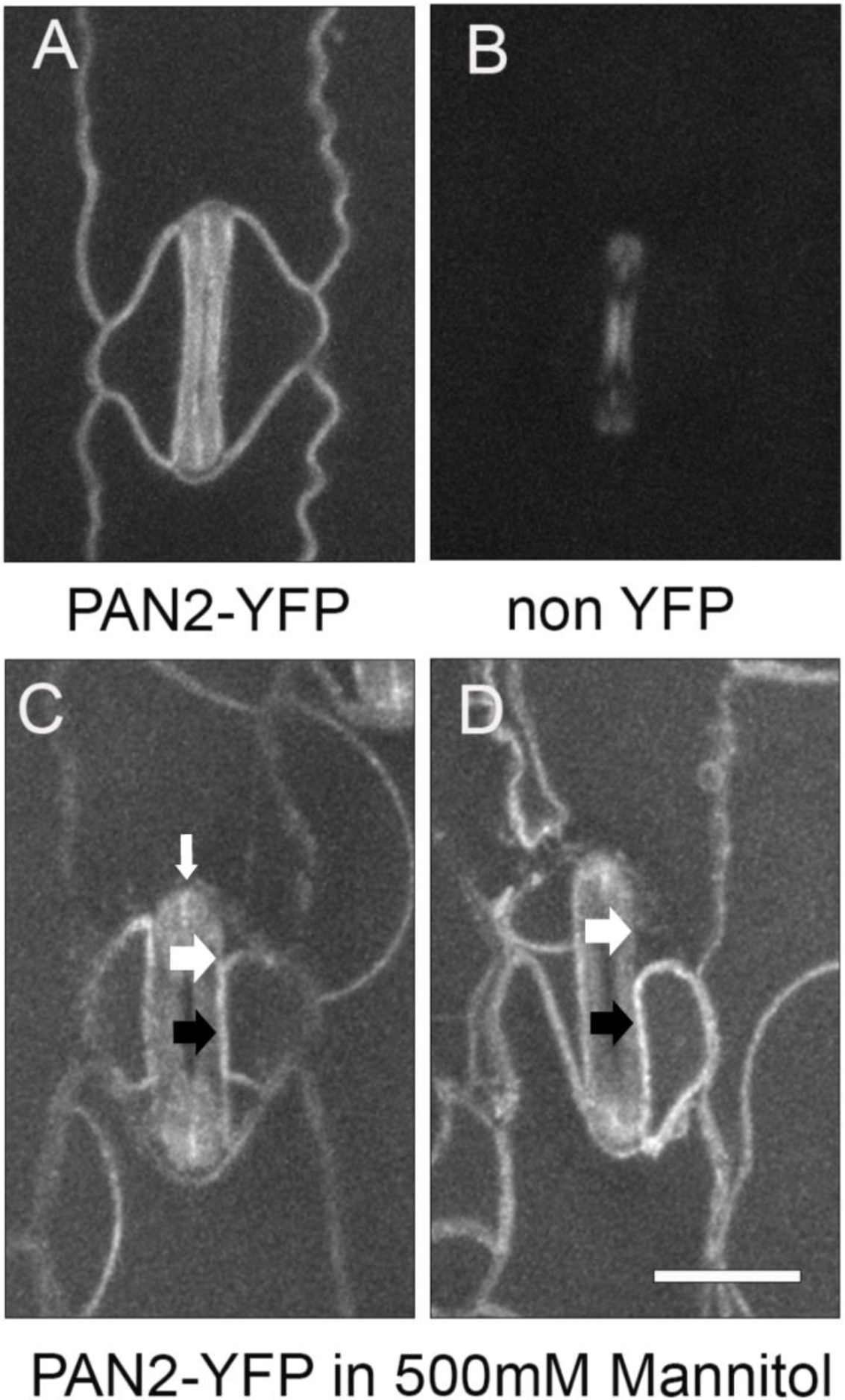
PAN2 is expressed in the subsidiary cells and pavement cells. (A, B) Fully expanded leaf 5 was examined in transgenic maize plants expressing PAN2-YFP (A) and a sibling plant without PAN2-YFP (B). Plants were imaged on the same day with the identical acquisition settings and images were scaled identically. Panel B shows guard cell autofluorescence. (C, D) Representative images of fully expanded leaf 4 tissue from plants expressing PAN2-YFP incubated in 500 mM mannitol for plasmolysis. Expression can be seen in the subsidiary cells. Black arrows indicate where the subsidiary cell plasma membrane is still adjacent to the guard cell. White arrows indicate where the subsidiary cell membrane has pulled away, indicating the membrane fluorescence in the subsidiary cell. Small vertical white arrow in A shows a bright line between the two guard cells, which may be PAN2-YFP or autofluorescence. Scale bar = 20µm, all images scaled identically.

**Fig. S8.**
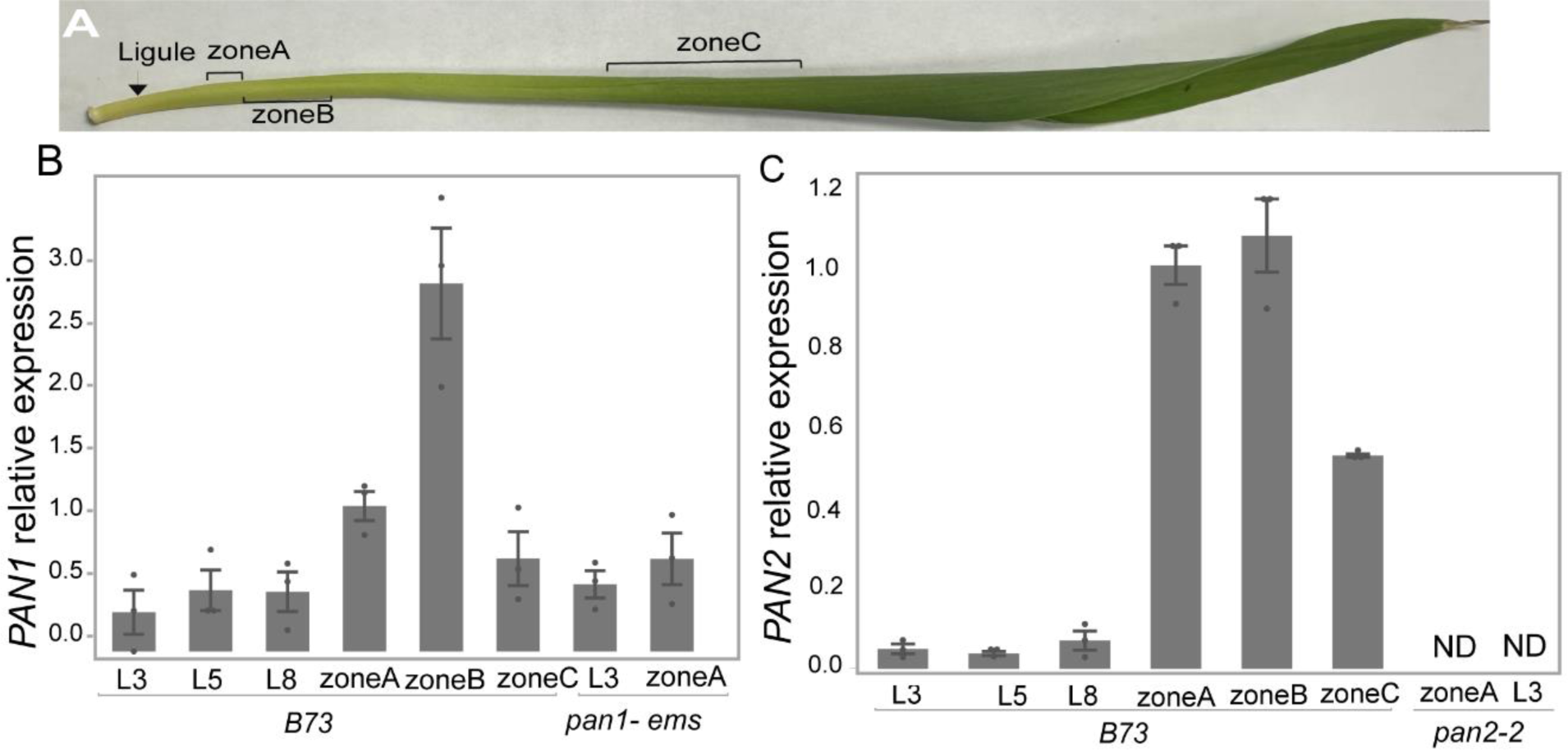
Relative expression levels of *PAN1* and *PAN2* in division zones and fully expanded Leaf 3, 5, and 8. (A) Diagram of developing leaf 5 showing the zones used for qRT-PCR analysis. (B) qRT-PCR analysis showing expression of PAN1 in fully expanded leaf 3 (L3), leaf 5 (L5), or leaf 8 (L8), or in leaf zones of unexpanded leaves as marked in panel A. Gene expression was measured in B73, and the *pan1-ems* null mutant was used as negative control (L3 and zone A only). (B) qRT-PCR analysis showing expression of PAN2. Labels are as in panel 1. The null mutant *pan2-2* was used as a negative control. The data are presented as mean ± SE of three biological replicates; each point represents each replicate’s relative expression value. The fold change of each gene expression is computed relative to ubiquitin, and normalized relative to zone A. In *pan2-2*, one or more samples had no determined (“ND”) expression. Exact CT values calculated relative values, and pairwise *t*-tests and *p*-values are listed Supplementary Tables S6 and S7.

