## Supplemental Figures for "Stomatal closure in maize is mediated by subsidiary cells and the PAN2 receptor"

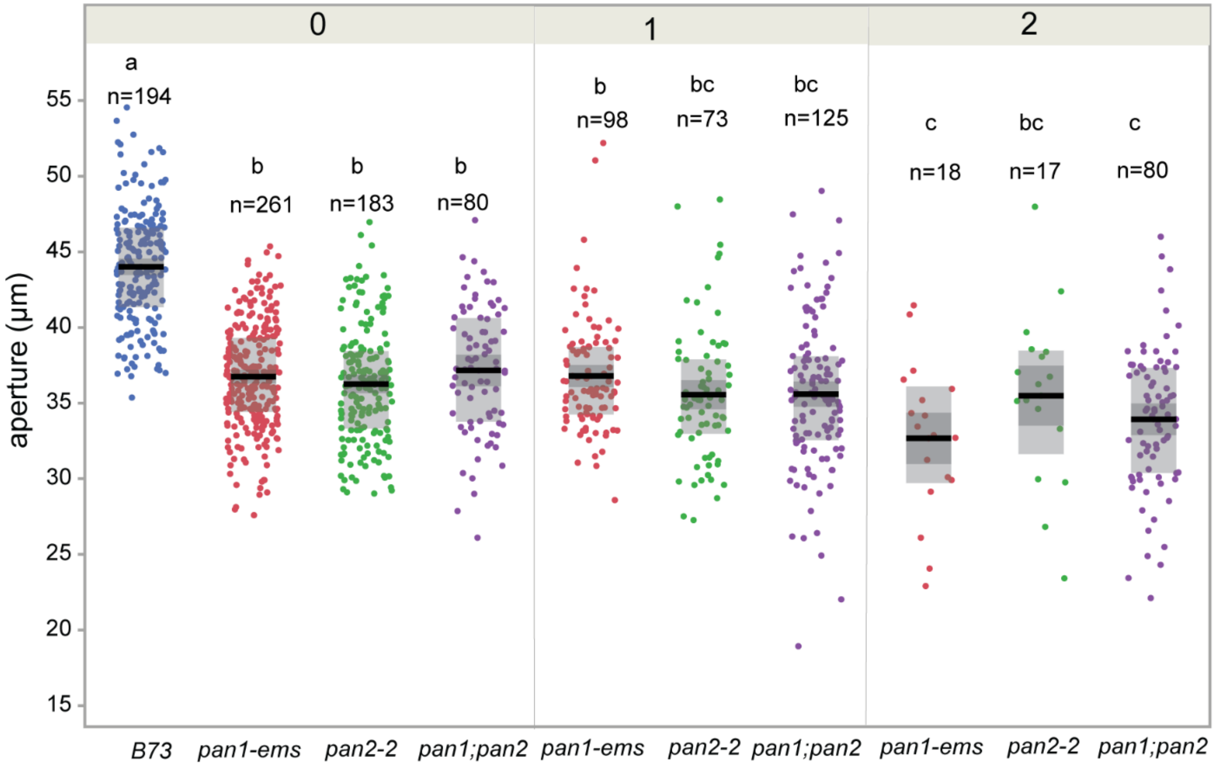


**Fig. S1 Guard cells in *pan1, pan2,* and *pan1;pan2* are shorter than those in B73, in leaf 5.** Guard cell length was measured using confocal images. Stomata were classified as having 0 abnormal (i.e., normal), 1 abnormal, or 2 abnormal subsidiary cells. Each data point represents a single guard cell pairs length, with 3-5 plants per genotype used. n represents the numbers of guard cell pairs used for measurement. Black bars indicate means; dark grey boxes are standard errors and larger light grey boxes indicate interquartile range. The number (0, 1, 2) above the panel represents the number of abnormal subsidiary cells in the stomate. Genotypes with different letters represent the guard cell lengths that are significantly different from each other ($p \leq 0.05$) Tukey Kramer Test.

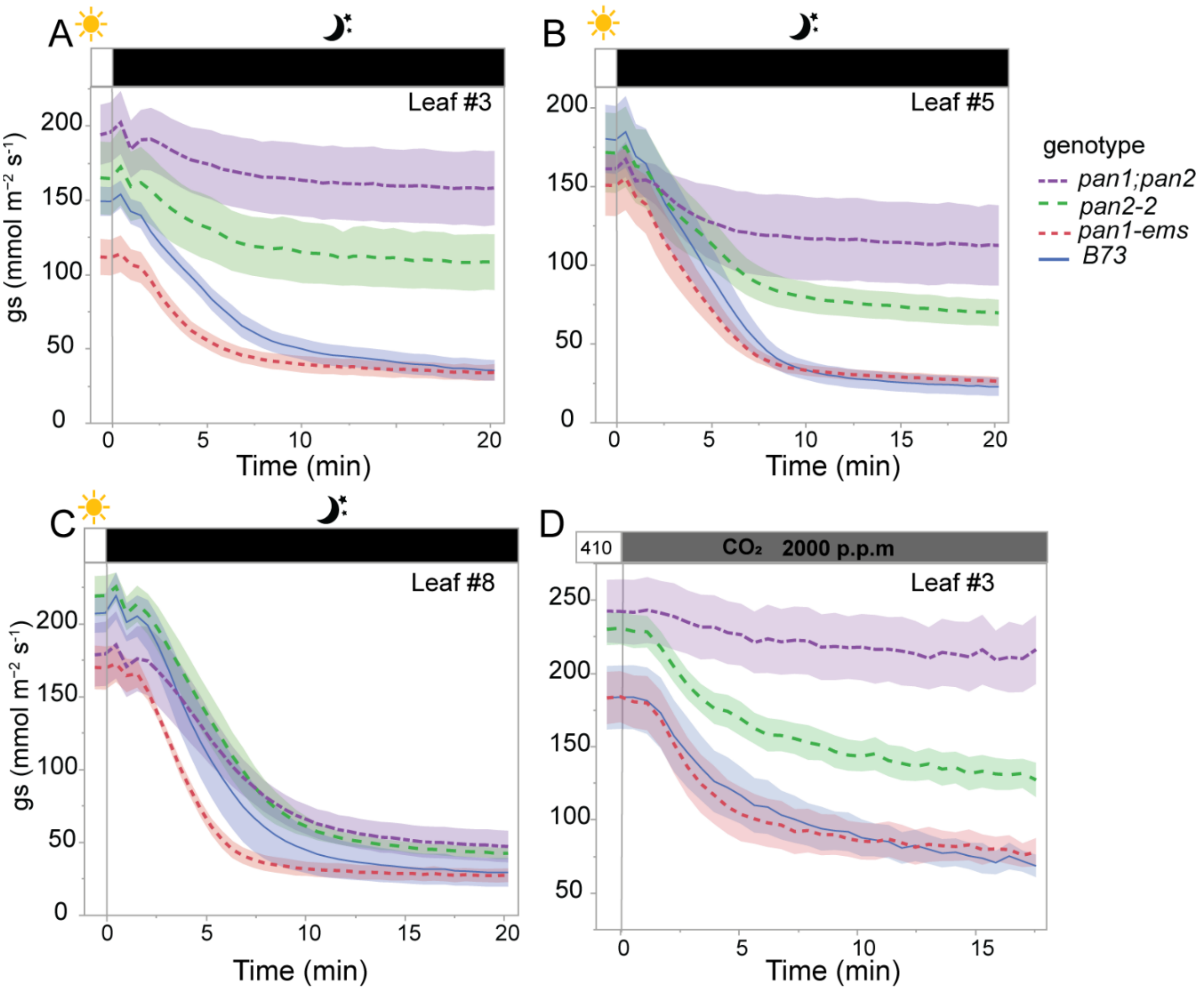


**Fig. S2 Stomatal closure is impaired in *pan2* and *pan1;pan2* and but is normal in *pan1* (absolute conductance)**. The normalized stomatal conductance data plotted in Figure 2 are plotted here as absolute stomatal conductance. Absolute stomatal conductance ($g_s$) of *pan2*, *pan2;pan*1, *pan1*, and B73 in response to darkness in leaf 3 (A), leaf 5, (B), or leaf 8 (C) was measured over time. PPFD was changed from 1000 to 0 at time zero. (D) Normalized stomatal conductance ($g_s$) in leaf 3 in response to elevated $CO_2$ was measured over time. The $CO_2$ concentration of from 410 ppm to 2000 ppm at zero time. Measurements were taken every 5 seconds. The data presented are the mean ± SE of at least 5 individual plants. Pertains to Figure 2.

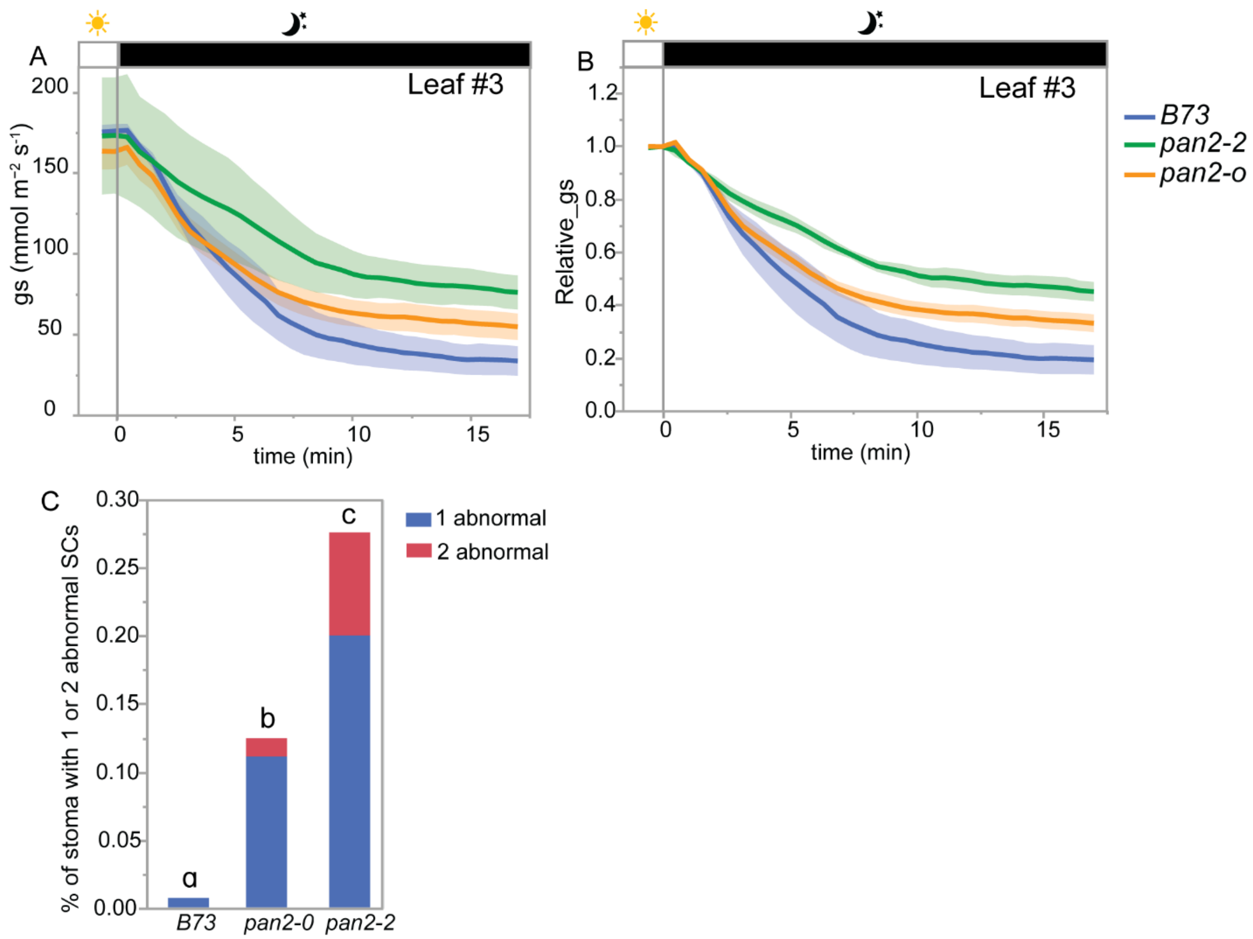


**Fig. S3 Comparison of stomatal developmental defects between *pan2-2* (strong allele) and *pan2-O* (weak allele).** (A) The $g_s$ of B73, *pan2-o*, and *pan2-2 was* measured in leaf 3 using in response to darkness. Light was turned off from 1000 to 0 at zero for the darkness response. The data presented are the mean ± SE of at least 3 individual plants. (B) Relative $g_s$ of B73, *pan2-o* and *pan2-2* computed for each measured plant by normalizing $g_s$ to the observed steady initial $g_s$ value. (C) Percentage of abnormal subsidiary cells (SCs) in B73, *pan2-O*, and *pan2-2* in leaf 3. All the plants were planted simultaneously. At least 300 stomata complexes were counted for each plant, and at least three plants were counted for each genotype. Genotypes with different letters represent the percentage of stomata with one and two abnormal subsidiary cells that are significantly different from each other (p≤0.05). Statistical analysis is performed based on the Student's t-test. Pertains to Figure 2.

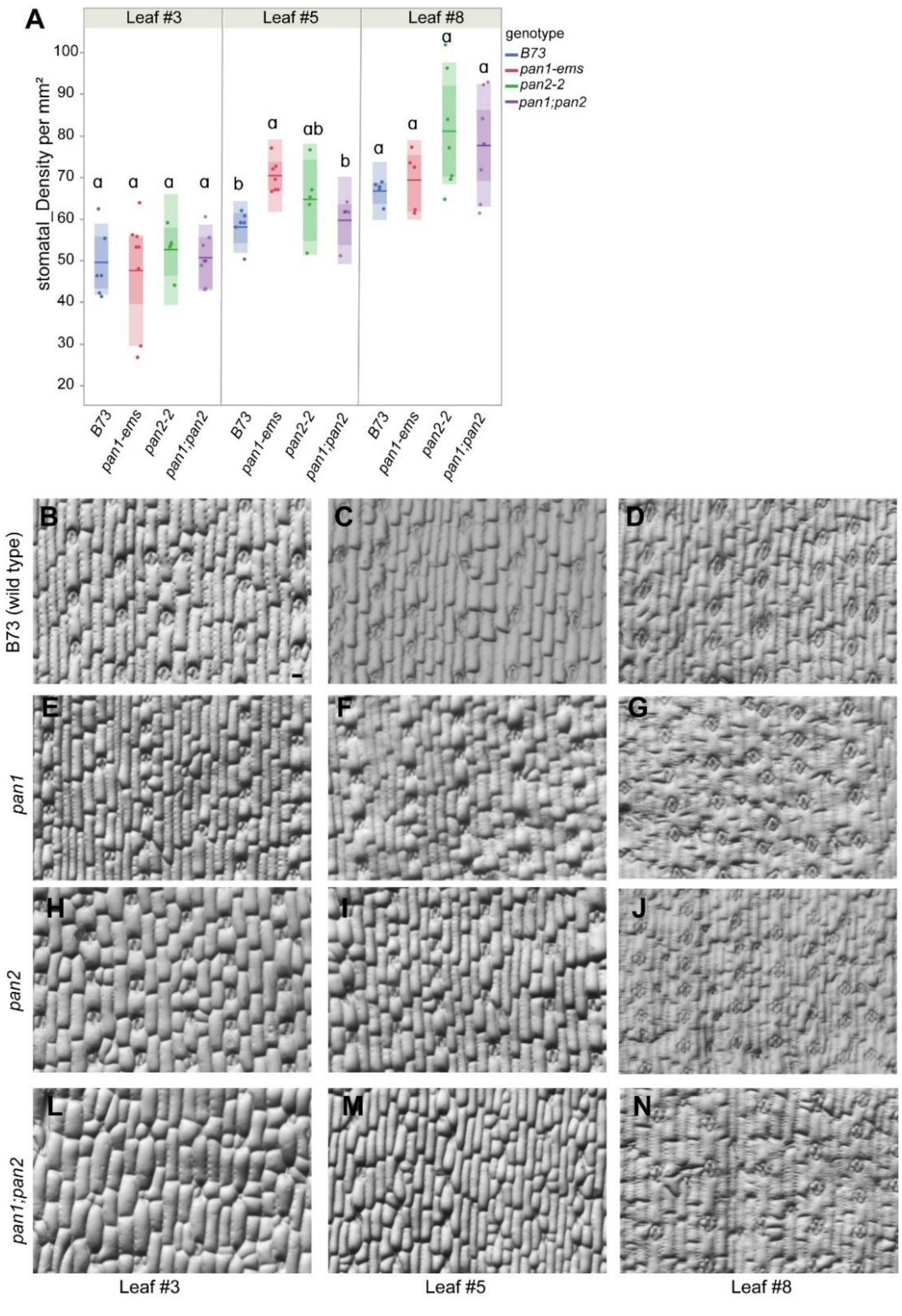


**Fig. S4 Stomatal density in B73, *pan1*, *pan2*, and *pan2;pan1*** (A) Stomatal density was measured using epidermal glue impressions. Density was measured in the same plants used for conductance measurements. Bars indicate means; dark shaded boxes indicate standard errors and lighter colored boxes indicate interquartile means. Genotypes with different letters indicate densities that are significantly different from each other ($p \leq 0.05$) based on Student's t-test. (B-N) The representative images of each genotype from leaf 3, 5, and 8. All the images were at the same scale. Scale bar in B is 50 μm. Pertains to Figure 2.

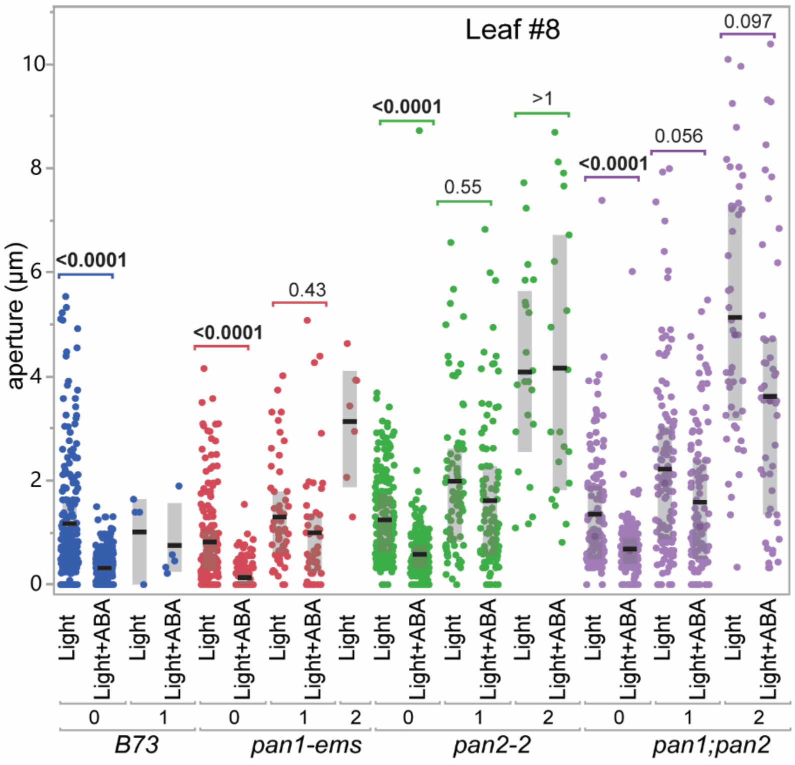


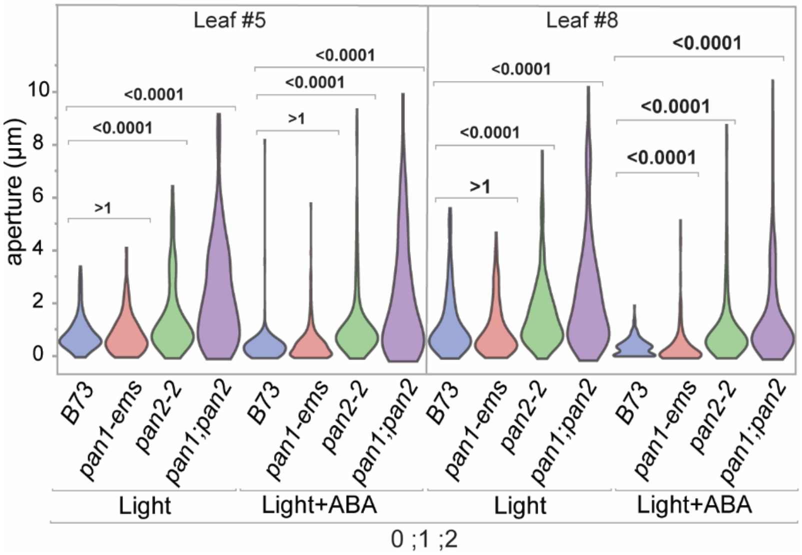


**Fig. S5 Stomatal aperture of different genotypes in response to ABA.** (A) Fully expanded leaf 8 segments from B73, *pan1,* or *pan1;pan2* was incubated in an opening buffer under 500 µm/m$^2$/s light, +/- 1 mM ABA. Stomata were classified as having 0, 1, or 2 abnormal subsidiary cells. Each data point represents a single stomatal complex, with 3-5 plants per genotype & treatment were used. Black bars indicate medians; grey boxes indicate interquartile ranges. (B) Violin plots of aperture measurements from all stomates with any number abnormal subsidiary cells, which represents the entire leaf sample (i.e., similar to gas exchange measurements). Leaf 5 is shown in the left panel; leaf 8 is shown on the right. Padj-values from a Bonferroni-corrected Wilcoxon comparison are as labeled. Pertains to Figure 3.

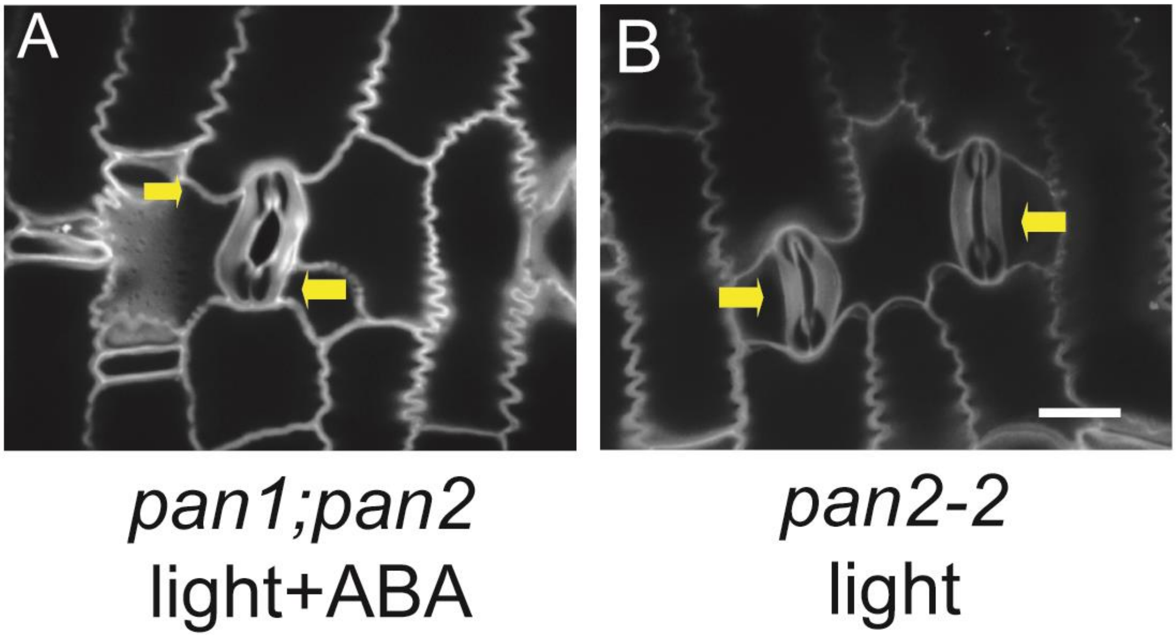


**Fig. S6 Representative images of "curved" guard cells from *pan2-2* and *pan1;pan2* mutants with or without treatment with ABA.** Leaf pieces of fully expanded leaves were treated in opening buffer in the presence or absence of 1 mM ABA. The yellow arrows indicate the direction of the subsidiary cell "push". The scale bar is 20 μm. Pertains to Figure 4.

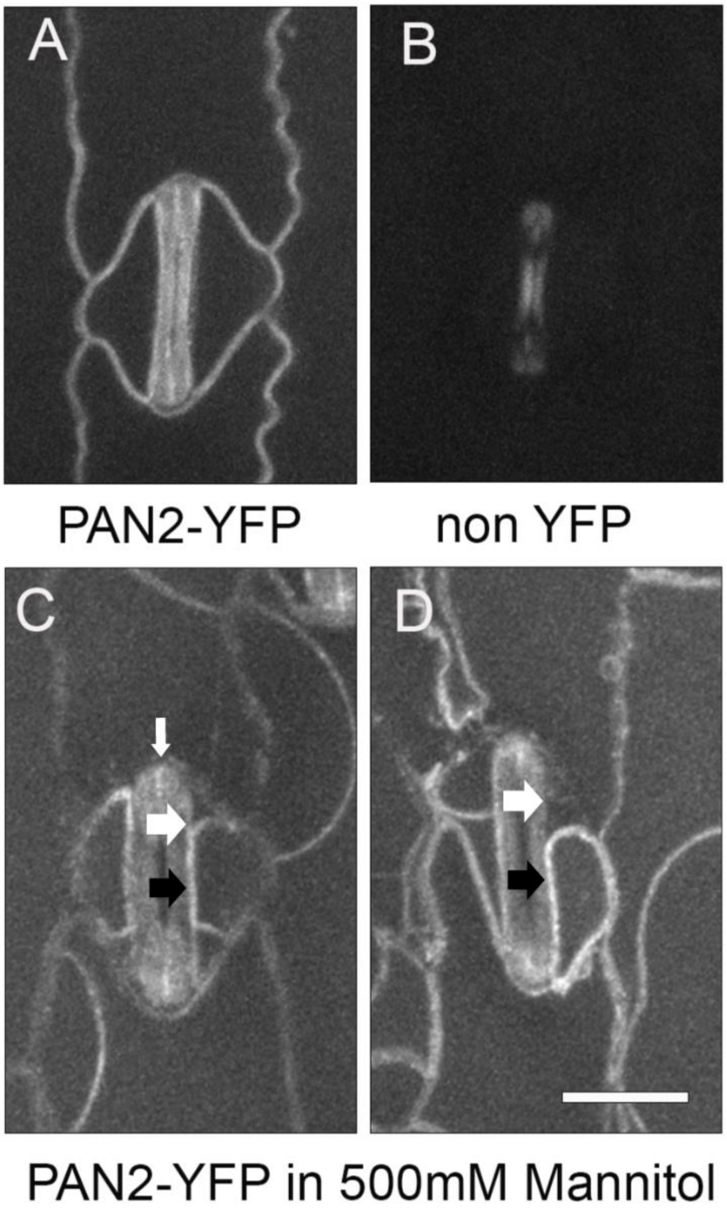


**Fig. S7 PAN2 is expressed in the subsidiary cells and pavement cells.** (A, B) Fully expanded leaf 5 was examined in transgenic maize plants expressing PAN2-YFP (A) and a sibling plant without PAN2-YFP (B). Plants were imaged on the same day with the identical acquisition settings and images were scaled identically. Panel B shows guard cell autofluorescence. (C, D) Representative images of fully expanded leaf 4 tissue from plants expressing PAN2-YFP incubated in 500 mM mannitol for plasmolysis. Expression can be seen in the subsidiary cells. Black arrows indicate where the subsidiary cell plasma membrane is still adjacent to the guard cell. White arrows indicate where the subsidiary cell membrane has pulled away, indicating the membrane fluorescence in the subsidiary cell. Small vertical white arrow in A shows a bright line between the two guard cells, which may be PAN2-YFP or autofluorescence. Scale bar = 20µm, all images scaled identically.

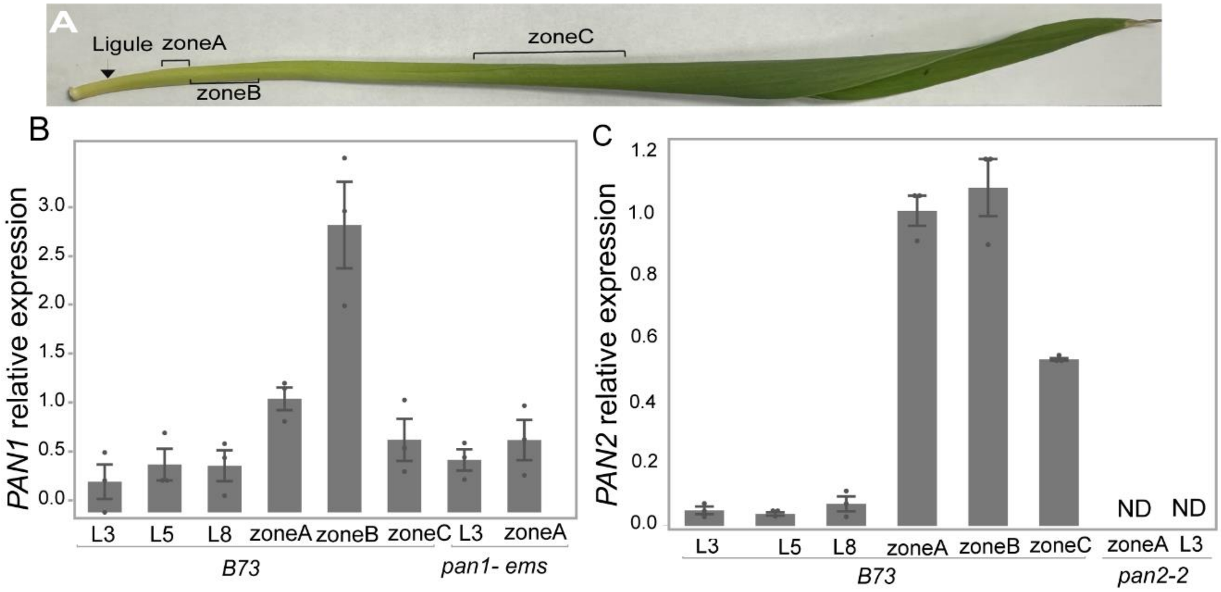


**Fig. S8 Relative expression levels of *PAN1* and *PAN2* in division zones and fully expanded Leaf 3, 5, and** 8. (A) Diagram of developing leaf 5 showing the zones used for qRT-PCR analysis. (B) qRT-PCR analysis showing expression of PAN1 in fully expanded leaf 3 (L3), leaf 5 (L5), or leaf 8 (L8), or in leaf zones of unexpanded leaves as marked in panel A. Gene expression was measured in B73, and the *pan1-ems* null mutant was used as negative control (L3 and zone A only). (B) qRT-PCR analysis showing expression of PAN2. Labels are as in panel 1. The null mutant *pan2-2* was used as a negative control. The data are presented as mean ± SE of three biological replicates; each point represents each replicate's relative expression value. The fold change of each gene expression is computed relative to ubiquitin, and normalized relative to zone A. In *pan2-2*, one or more samples had no determined (“ND”) expression. Exact CT values calculated relative values, and pairwise *t*-tests and *p*-values are listed Supplementary Tables S6 and S7.
